# 3D Reconstruction of Murine Mitochondria Exhibits Changes in Structure Across Aging Linked to the MICOS Complex

**DOI:** 10.1101/2022.03.22.485341

**Authors:** Zer Vue, Edgar Garza-Lopez, Kit Neikirk, Prasanna Katti, Larry Vang, Heather Beasley, Jianqiang Shao, Andrea G. Marshall, Amber Crabtree, Alexandria C. Murphy, Brenita C. Jenkins, Praveena Prasad, Chantell Evans, Brittany Taylor, Margaret Mungai, Mason Killion, Dominique Stephens, Trace A. Christensen, Jacob Lam, Benjamin Rodriguez, Mark A. Phillips, Nastaran Daneshgar, Ho-Jin Koh, Alice Koh, Jamaine Davis, Nina Devine, Saleem Muhammod, Estevão Scudese, Kenneth Ryan Arnold, Valeria Vanessa Chavarin, Ryan Daniel Robinson, Moumita Chakraborty, Jennifer A. Gaddy, Mariya Sweetwyne, Genesis Wilson, Elma Zaganjor, James Kezos, Cristiana Dondi, Anilkumar K. Reddy, Brian Glancy, Annet Kirabo, Anita M. Quintana, Dao-Fu Dai, Karen Ocorr, Sandra A. Murray, Steven M. Damo, Vernat Exil, Blake Riggs, Bret C. Mobley, Jose A. Gomez, Melanie R. McReynolds, Antentor Hinton

**Author notes:** Corresponding Author: Antentor Hinton, Department of Molecular Physiology and Biophysics Vanderbilt University. These authors share co-first authorship (Zer Vue, Edgar Garza-Lopez, Kit Neikirk, Prasanna Katti). These authors share senior authorship (Melanie R. McReynolds, Antentor Hinton).

## Abstract

**Background:** During aging, muscle gradually undergoes loss of function including sarcopenia, losing mass, strength, endurance, and oxidative capacity. While mitochondrial aging is associated with decreased mitochondrial capacity, the genes associated with morphological changes in mitochondria during aging still require further elucidation. Furthermore, it is not completely understood how 3D mitochondrial structures are altered during aging in skeletal muscle and cardiac tissues.

**Methods:** We measured changes in mitochondrial morphology and mitochondrial complexity during the aging of murine gastrocnemius, soleus, and cardiac tissues using serial block face-scanning electron microscopy and 3D reconstruction. Lipidomic and metabolomic analysis elucidated concomitant changes associated with aging. We also used qPCR, transmission electron microscopy quantification, Seahorse Analyzer, and metabolomics to evaluate changes in mitochondria morphology and function upon loss of the MICOS complex.

**Results:** We identified significant changes in 3D mitochondrial size and network configuration in murine gastrocnemius, soleus, and cardiac tissue during aging. These changes were concomitant with loss of mitochondria contact site and cristae organizing system (MICOS) gene expression during aging. Mitochondrial morphology was similar between aged mice and young mice. We show an age-related loss of the MICOS complex (Chchd3, chchd6, and Mitofilin) while their knockout results in alterations in mitochondrial morphology. Given the critical role of mitochondria in maintaining cellular metabolism, we perform cellular metabolic profiling of young and aged tissues. Metabolomics and lipidomics showed profound alterations, including in membrane integrity, that support our observations of age-related changes in these muscle tissues.

**Discussion:** In tandem, our data suggest a relationship between the MICOS complex and aging, which could be linked to disease states with further 3D reconstruction studies. Our study highlights the importance of understanding tissue-dependent 3D mitochondrial phenotypical changes which occur across aging with evolutionary conservation between *Drosophila* and murine models.

**Graphical Abstract:** 

## INTRODUCTION

Sarcopenia is the loss of muscle mass associated with aging, which impacts the quality of life. Sarcopenia predominately affects type II muscle fibers, but also affects type I fibers.

Mitochondrial dysfunction is a hallmark of aging^1^. With age, sarcopenia may occur in skeletal muscle, leading to a body mass-independent loss of skeletal function^2^. Decreased expression of genes associated with mitochondrial dynamics and loss of function is known to contribute to sarcopenia and other age-related diseases ^3^. A hallmark of aging is alterations in mitochondria structure ^1^. Thus, mitochondria are a key target for the development of potential therapeutics for age-related pathologies ^1,3^. Mitochondria maintain finely regulated structures, through various mechanisms, including fission and fusion ^4^. Because their structure changes dynamically as mitochondria shift from fission to fusion, many important mitochondrial functions depend on their structure^5–7;^ therefore, it is important to understand mitochondrial structural changes over time. Essential mitochondrial functions are associated with the cristae, the inner folds of the mitochondrial membrane, which house the oxidative phosphorylation machinery^6^ and various transporters. Our objective was to determine how mitochondrial structures change during aging. We hypothesized that age-related alterations in metabolism and lipids cause increased mitochondrial fragmentation over time with concomitant losses in mitochondrial cristae integrity.

The ultrastructure and phenotypical variation of mitochondria play important roles in mitochondrial function. Disruption of optic atrophy 1 (OPA-1), an inner membrane protein that regulates mitochondrial fusion, causes mitochondrial fragmentation and affects the dimensions, shapes, and sizes of the cristae^6^. Furthermore, disruption of dynamin-related protein-1 (DRP1), a protein associated with mitochondrial fission, causes elongated mitochondria and resistance to cristae remodeling^8,9^. Nanotunnels, or “mitochondria-on-a-string”, are thin, double-membrane protrusions that allow mitochondria to communicate across distances. Nanotunnels may increase in mitochondrial disease^10,11^ and may be associated with mitochondrial dysfunction during aging. Thus, the alterations in mitochondrial structure and bioenergetics may simultaneously drive pathologies.

Mutations in genes that regulate the morphology of cristae are associated with aging cardiomyocytes^12^. These proteins are located at the crista junctions in the inner membrane and are a part of the mitochondrial contact site and cristae organizing system (MICOS) complex, which is important for maintaining mitochondrial shape and size^13^. DRP1 or OPA-1 loss can similarly affect mitochondria morphology^14,15^. Cristae membranes contain the electron transport chain complexes and ATP synthases for oxidative phosphorylation^16–18^. Since mitochondrial morphology affects function, altering the structure by knocking out MICOS-associated genes or OPA-1, a GTPase, could alter the metabolism and function of mitochondria during aging^16–18^. We sought to better understand how the MICOS complex affects gross mitochondrial structure beyond only cristae, as well as how mitochondrial structure changes in aging. Critically, we predicted that MICOS-associated gene products are lost during aging; therefore, loss of MICOS-associated genes would mimic the change in mitochondria morphology observed in aging. Mitochondria have a tissue-dependent response to the environment ^19^, which may be due to heterogeneity in mitochondrial DNA (mtDNA) quality check mechanisms across tissues ^20^.

Using manual contour tracings, 3D reconstruction is an important tool to understand specific phenotypes of mitochondria as well as how they may differ across tissue types. Therefore, we sought to better understand the alterations of mitochondrial structure in aging; we compared size, shape, quantity, complexity, and branching of mitochondria using 3D reconstructions of aged gastrocnemius, soleus, and cardiac tissue in 3-month and 2-year mice. We observed an age-related loss of the MICOS complex in both qPCR and western blotting. Beyond this, we used CRISPR/Cas9 on myotubes to knockout (KO) three genes of the MICOS complex: *Chchd3* (Mic19), *Chchd6* (Mic25), and *Mitofilin* (Mic60) to understand how the MICOS complex may present phenotypically similar to the aging process through modulation of mitochondrial size, morphology, and oxygen consumption rate. To further understand factors that may affect mitochondrial structure in aging, multivariate analysis was used to identify changes in metabolites and characterize the lipidome across aging to understand commonalities to metabolic changes with loss of the MICOS complex. Finally, a *Drosophila* model was used to further explore the role of the MICOS complex beyond murine and to elucidate changes in heart function and MICOS gene expression in aging.

## RESULTS

### Aging Results in Smaller Mitochondria in Murine Gastrocnemius, Soleus, and Cardiac Tissue

The gastrocnemius is a mixed muscle with both oxidative fibers, containing many mitochondria, and glycolytic fibers, with few mitochondria ^21^. This heterogeneity makes the gastrocnemius ideal to study changes in mitochondrial dynamics. In contrast, soleus tissue is made up predominately of slow-twitch muscle fibers, reliant on oxidative metabolism dependent on the relatively increased mitochondrial content ^22^. Mitochondria in cardiac tissue, as we have previously studied ^23^, further have differential roles more reliant on efficient energy transfer to myofibrils and constant ATP production to maintain contractile function ^24^. Since many mitochondrial functions depend on structure ^5–7^, it is important to examine how mitochondrial structure changes over time. We hypothesized that increasing mitochondrial fragmentation over time is concomitant with losses in mitochondrial crista integrity.

Our objective was to determine how aging alters mitochondrial networks and individual mitochondrial structures. We imaged gastrocnemius, soleus, and cardiac biopsies from adolescent (3-month-old) and aged (2-year-old) mice by serial block-face scanning electron microscopy (SBF-SEM) with resolutions of 10 nm for the x- and y- planes and 50 nm for the z- plane, which allows visualization of the electron connectome. Approximately 50 intermyofibrillar (IMF) mitochondria were segmented from each image stack (Figure 1A-F) and a 3D surface view was generated (Figure 1A’-F’). We analyzed IMF, as opposed to other mitochondrial subpopulations such as subsarcolemmal, as IMF mitochondria are larger and displayed more significant age-related changes. We analyzed mitochondria sub-network volumes from four ROIs with an average of 175 mitochondria for each mouse (n = 3), for a total of 500 mitochondria. Mitochondrial networks in aged skeletal muscle tissue mice showed largely interconnected mitochondria that were not composed of a single reticulum (Figures 1A’’–D’’).

**Figure 1:**
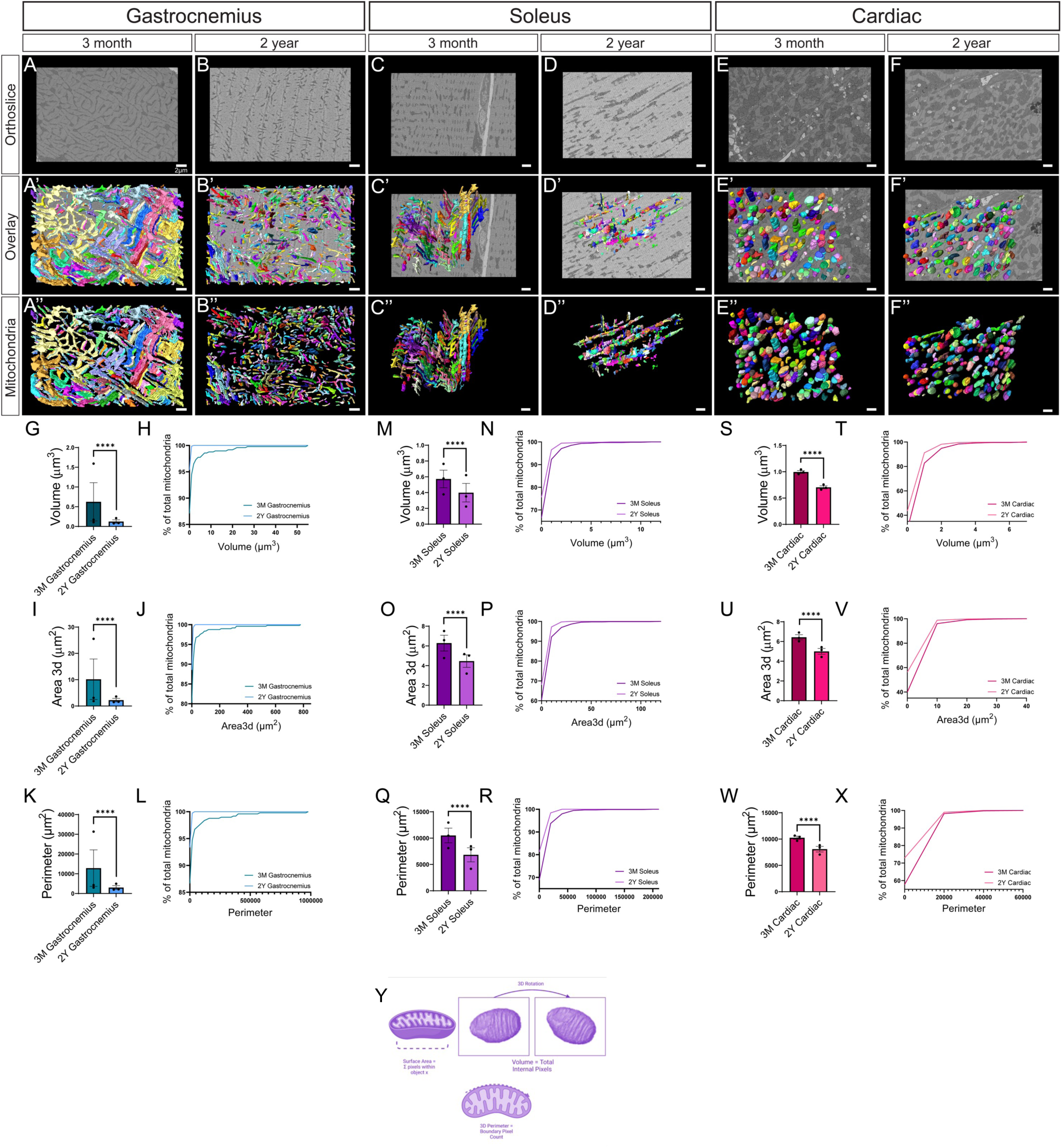
Decreased mitochondrial size and volume in gastrocnemius, soleus, and cardiac muscle of aged mice in SBF-SEM 3D reconstructions. **(A-B)** Representative SBF-SEM orthoslice for male murine gastrocnemius, **(C-D)** soleus, and (E-F) cardiac tissue. (A’-B’) 3D reconstructions of mitochondria (various colors) in gastrocnemius, (C’-D’) soleus, and (E’-F’) cardiac tissue of three-month and 2-year mice overlaid on ortho slices. (A’’-B’’) Pseudo-colored individual mitochondria in gastrocnemius, (C’’-D’’) soleus, and (E’’-F’’) cardiac tissue differentiate micro-level changes. **(G-X)** Quantification of 3D reconstructions, with each dot representing the overall average of all mitochondria quantified for each mouse. **(G)** Mitochondrial volume in gastrocnemius muscle of 3-months and 2-year murine samples, and (H) mitochondrial volume distributed on basis of percent of total mitochondria to visualize relative heterogeneity. **(I)** Mitochondrial 3D area in gastrocnemius muscle of 3-months and 2-year murine samples, and **(J)** mitochondrial area distributed on basis of percent of total mitochondria to visualize relative heterogeneity. **(K)** Mitochondrial perimeter in gastrocnemius muscle of 3-months and 2-year murine samples, and **(L)** mitochondrial perimeter distributed on basis of percent of total mitochondria to visualize relative heterogeneity. These same quantifications are also displayed in **(M-R)** soleus and **(S-X)** cardiac tissues. Cartoon representations of metrics to calculate **(Y)** mitochondrial volume, **(Z)** perimeter, and (AA) perimeter. Approximately 550 mitochondria analyzed for each tissue type and age cohort (n=3 mice per age cohort). Significance values **** represents *p* ≤ 0.0001.

As in our previous study, we found cardiac tissue mitochondria remained relatively clumped with no apparent distribution changes (Figure 1E”-F”). We found that across all tissue types, volume, area, and perimeter of mitochondria of samples from 2-year-old mice were significantly lower than those from 3-month-old mice (Figures 1G-X). Here mitochondria volume is a measure of their total capacity (Figure 1Y), while area represents a measure analogous to surface area (Figure 1Z) and perimeter represents boundary pixel count (Figure 1AA).

While there was some variability across the three mice for each age cohort (Supplemental Figure 1), this heterogeneity is more pronounced in the gastrocnemius muscle (Figure 1G-H). Gastrocnemius also showed a greater reduction in mitochondrial size and surface area, demonstrating much smaller mitochondria no longer presenting as hyperbranched. Soleus tissue similarly demonstrated heterogeneity between the samples with similar reductions in mitochondrial volume across aging (Figure 1M-R). In contrast to skeletal muscle, cardiac tissue shows a much more homogenous mitochondrial population (Figure 1S-X). In general, the trend showed a downward trajectory emblematic of increased fragmentation and smaller mitochondria. This showed that the size and length of mitochondria change with age, so we sought to further elucidate the complexity of mitochondria, which is implicated in mitochondria communication.

### Aging Causes Altered Mitochondrial Phenotypes with Poorly Connected Mitochondria with Decreased Branching in Murine Gastrocnemius, Soleus, and Cardiac Tissue

Next, we measured mitochondrial complexity to better understand changes in mitochondrial shape during aging. We hypothesized that fewer networks and simpler shapes would occur as aging and dysfunction continued. Since we observed in 3D reconstructions that mitochondrial populations are heterogeneous and diverse, we used mito-otyping, which is the karyotyping-like method for arranging mitochondria^25^, to capture the diversity of IMF mitochondria (Figure 2A- C). This analysis suggested, at every volumetric measurement, that mitochondria were smaller and less complex with age. Soleus and gastrocnemius tissue both showed branched distributions in the adolescent populations while aged populations appeared much less branched, in addition to having smaller volumes. Notably, skeletal muscle showed large phenotypical changes with aging while cardiac tissue does not show the same changes. To validate these changes, we utilized previously established methods were used to determine 3D mitochondrial complexity from 3D form factor measurements^25,26^. We measured mitochondrial complexity index (MCI) and sphericity to further understand changes in complexity (Figure 2D-O). MCI and sphericity are analogous measures to understand the general roundness of mitochondria (Figure 2P-Q). In gastrocnemius tissue, there was an increase in MCI concomitant with an increase in sphericity with aging (Figure 2D-G). This shows that, contrary to the appearance of mito-otyping, gastrocnemius exhibits increased sphericity of mitochondria. When examining the specific three male mice sampled for each age cohort, some variation exists for both metrics. Soleus tissue, which shows a similar heterogeneity, is exposed to be less complex and more spherical across aging (Figure 2H-K) although this is not as significant of a difference as that exhibited in gastrocnemius. Finally, cardiac tissue only shows a significant increase in MCI (Figure 2L-O).

**Figure 2:**
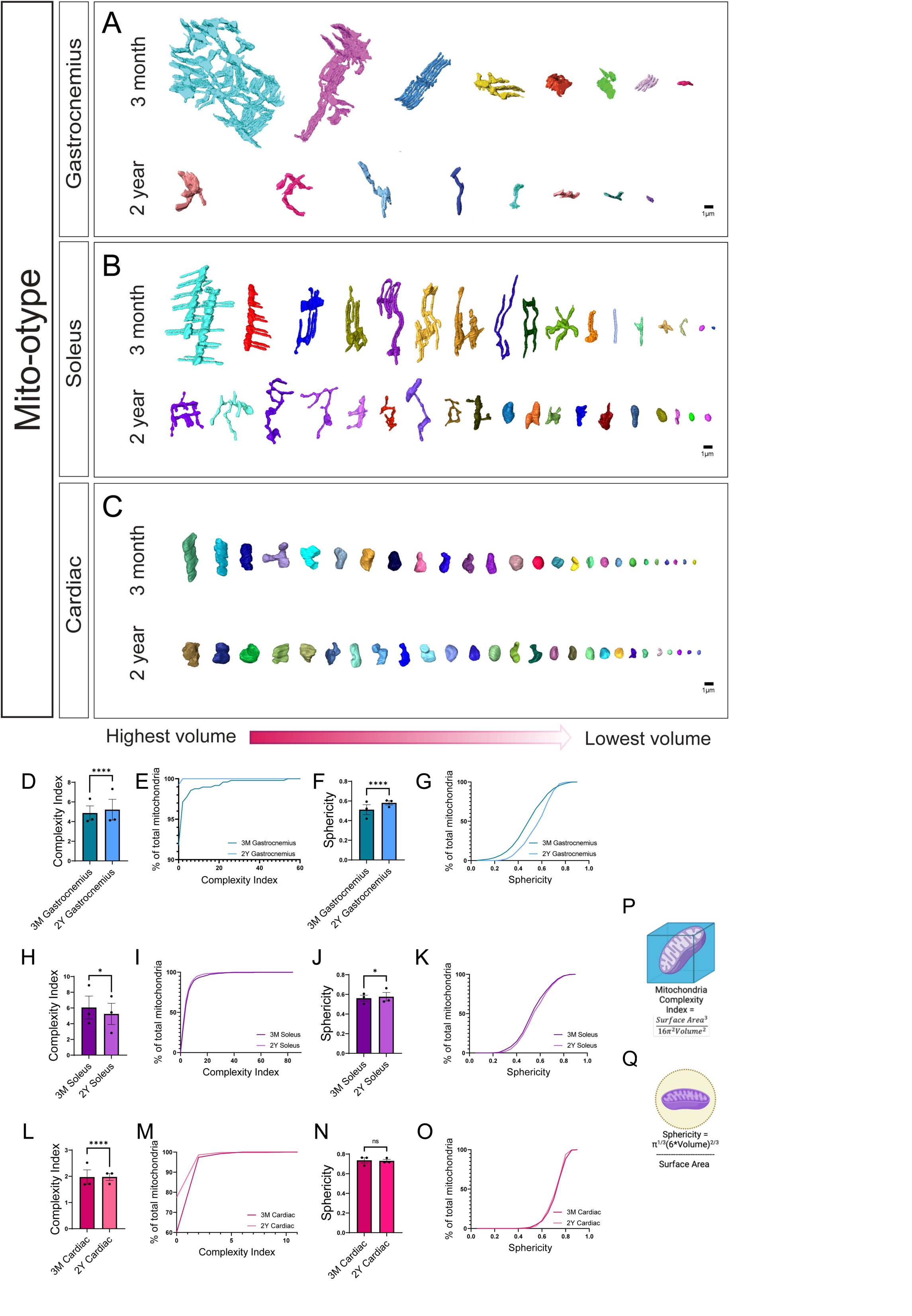
SBF-SEM 3D reconstruction in gastrocnemius, soleus, and cardiac muscle of aged mice showing altered mitochondrial networks. Representative examples of 3D reconstruction of mitochondria in (**A**) gastrocnemius, (**B**) soleus, and (**C**) cardiac tissue of three-month and 2-year mice ages organized by volume to show phenotypes of mitochondria. (**D**) Mitochondrial complexity index (MCI), a measure analogous to sphericity, in gastrocnemius muscle of 3-months and 2-year murine samples, and (**E**) MCI distributed on basis of percent of total mitochondria to visualize relative heterogeneity. (**F**) Sphericity in gastrocnemius muscle of 3-months and 2-year murine samples, and (**G**) mitochondrial sphericity distributed on basis of percent of total mitochondria to visualize relative heterogeneity. These same quantifications are also displayed in (**H-K**) soleus and (**L-O**) cardiac tissues. Cartoon representations of metrics to calculate (**P**) MCI and (**Q**) sphericity. Approximately 550 mitochondria analyzed for each tissue type and age cohort (n=3 mice per age cohort). Significance values * p≤ 0.05, **** *p* ≤ 0.0001.

Together, these data suggest that complexity is altered with age, but the specific way in which complexity changes is dependent on tissue type. These findings caused us to look at the MICOS complex to elucidate the role it has on mitochondrial structure and function, as a potential effector of age-related changes in mitochondrial structure.

### Age-Related Changes in the MICOS Complex Expression

The MICOS complex and OPA-1 are key players in mitochondrial biogenesis ^13,17,27^, but how their interactions regulate aging and mitochondrial structures is poorly understood. Since we observed mitochondrial dynamics change with aging, we measured transcripts for *Opa1*, *Chchd3* (*Mic19*), *Chchd6* (*Mic25*), and *Mitofilin* (*Mic60*), the genes for the principal MICOS complex subunits. We used reverse-transcriptase qPCR (RT-qPCR) to determine the effect of aging on transcription. Aging causes mitochondrial fragmentation ^1^, which is associated with the loss of *Opa1* and MICOS complex proteins ^17,28,29^. We hypothesized that aging would also lead to decreases in the associated MICOS complex proteins. We also sought to confirm prior studies that found *Opa1* loss across aging, by using *Opa1* as a positive control ^17,30–32^. We found a significant loss of both *Opa1* and MICOS complex subunits, as measured by loss of transcripts for the genes in skeletal muscle when comparing 3-month and 2-year samples (Figure 3A–D).

**Figure 3:**
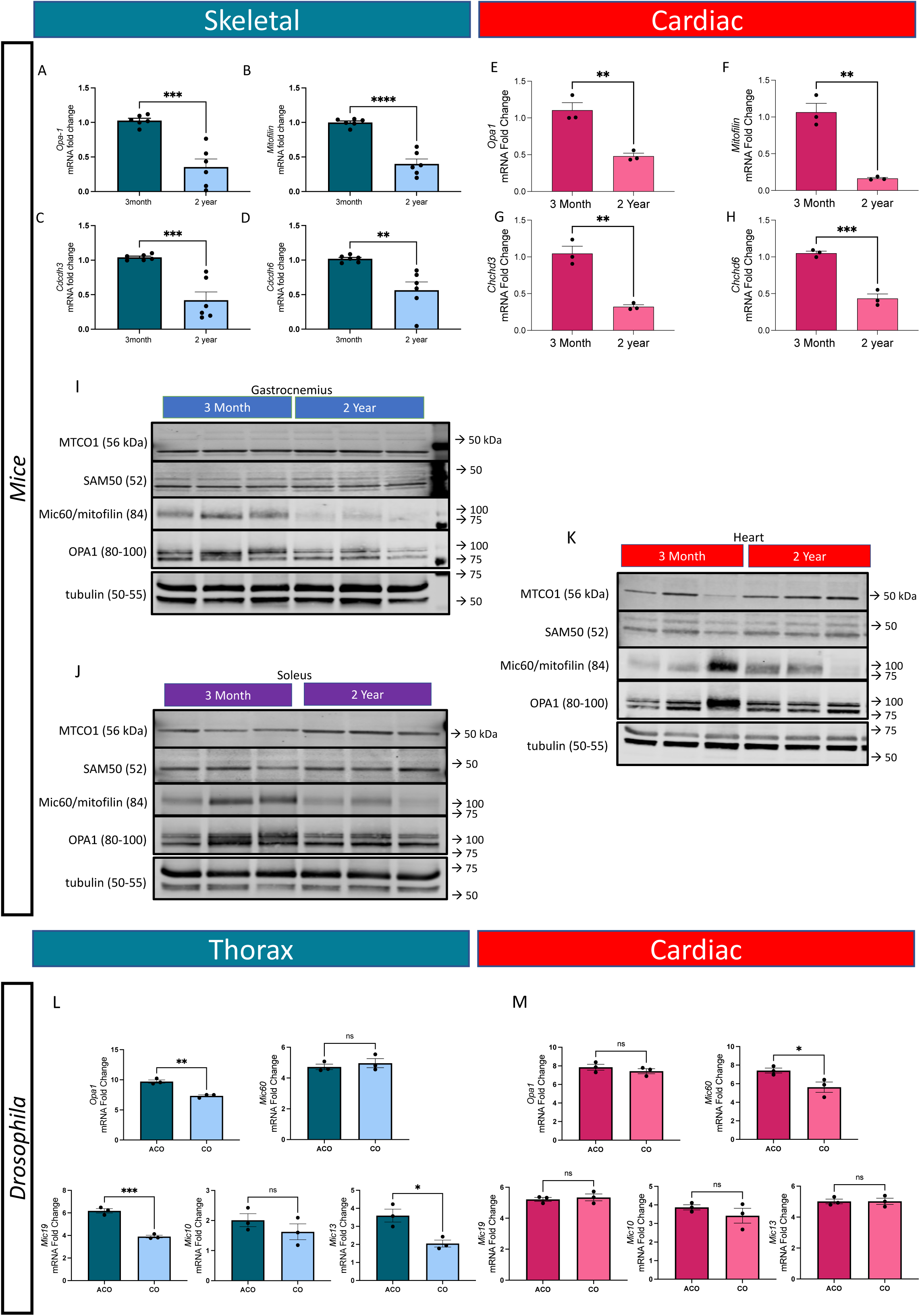
Changes in Expression of *Opa1* and MICOS genes in Aging Mouse Tissue and *Drosophila*. (**A-D**) Fold changes in the amount of *Opa1* and MICOS gene transcripts in mitochondria of skeletal muscle of 3-month-old and 2-year-old mice as measured by RT-qPCR. (**A**) *Opa1*, (**B**) *Mitofilin*, (**C**) *Chchd3*, and (**D**) *Chchd6* transcripts in skeletal muscle. **(E-H)** qPCR analyzing the gene transcript fold changes of Opa-1 and MICOS across aging in cardiac tissue. (**E**) *Opa1*, (**F**) *Mitofilin*, (**G**) *Chchd3*, (**H**) and *Chchd6* transcripts. (**I**) Altered protein expressions of MTCO1, SAM50, Mitofilin, OPA1, and tubulin in gastrocnemius, (**J**) soleus, and (**K**) cardiac tissue. (**L-M**) Altered *Drosophila* mitochondrial genes with age. (**L**) qPCR of *OPA1* and MICOS complex components in aging Drosophila in thorax and (**M**) cardiac tissue. Significance values * *p* ≤ 0.05, ** *p* ≤ 0.01, *** *p* ≤ 0.001, **** *p* ≤ 0.0001. For all qPCR experiments, n=6.

When looking at cardiac tissue (Figure 3E-H) we noticed a similar age-related loss of all MICOS genes products. These data suggest a correlation between mitochondrial morphological changes and decreases in expression of *Opa1* and MICOS subunits *Chchd3*, *Chchd6*, and *Mitofilin* during aging; however, they are not sufficient to demonstrate causation, therefore we did additional measurements to test if reduced protein expression levels may be responsible for the morphological changes. When looking at the MTCO1 (Cox-1) or Sam50 levels, there were not significant alterations across the aging process (Figure 3I-K). However, both gastrocnemius and skeletal tissue showed reduced protein levels of Mitofilin and OPA1 (Figure 3I-J); notably, however, there was a differential aging response with no age-related changes in protein levels in cardiac tissue (Figure 3K).

From there we sought to validate these results in another model. Using qPCR, we looked at differences in gene expression for MICOS complex as well as mitochondrial endoplasmic reticulum contact (MERC) genes between experimentally evolved Drosophila populations subjected to hundreds of generations for accelerated aging and their controls ^33^. We looked at MERC genes as the relationship between the MICOS complex and MERCs remains unclear, and we believed there may be concomitant loss. Specifically, we compared patterns between 21-day-old flies, with the accelerated aging flies displaying phenotypes of old age, where past studies have shown there are large differences in age-specific mortality, gene expression profiles, and metabolomic profiles ^33–35^. Flight muscle tissues, ATF-4, Drp, Marf, Opa1, MINOS1 (mic10), CHCHD3/6(mic19), QIL1(mic13), APOO (mic16/27), IP3R/Itpr, VDAC/Porin, Bip(grp78), GRP-75/Hsc70-5, and Ire1 all showed downregulated expression in indirect flight muscle of aged flies when compared with controls (Table 1; Figure 3L). However, only the expression changes associated with Drp, Marf, Opa1, CHCHD3/6, QIL1, APOO, IP3R, VDAC, and Bip were significant (p-value < 0.05; Unpaired T-test) (Table 1; Figure 3L). We saw a slight upregulated expression of ATF-6 and Mitofilin in the aged flies, however, these changes were not significant. ATF-4, Opa1, MINOS1, and Mitofilin all showed downregulated expression in the cardiac tissues of the aged flies when compared with controls, however, only the change in ATF-4 and Mitofilin expression were significant (Table 2; Figure 3M). In contrast, there was a significant increase in FGF-21 expression in the aged flies (Table 2). These validate that mitochondrial, MICOS, and MERC proteins are differentially impacted in cardiac and skeletal muscle tissue. Interestingly, together these also suggest that in some models MICOS complex has a greater dependence on the aging process.

**Table 1:**
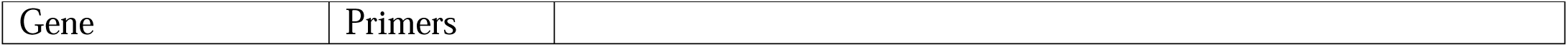
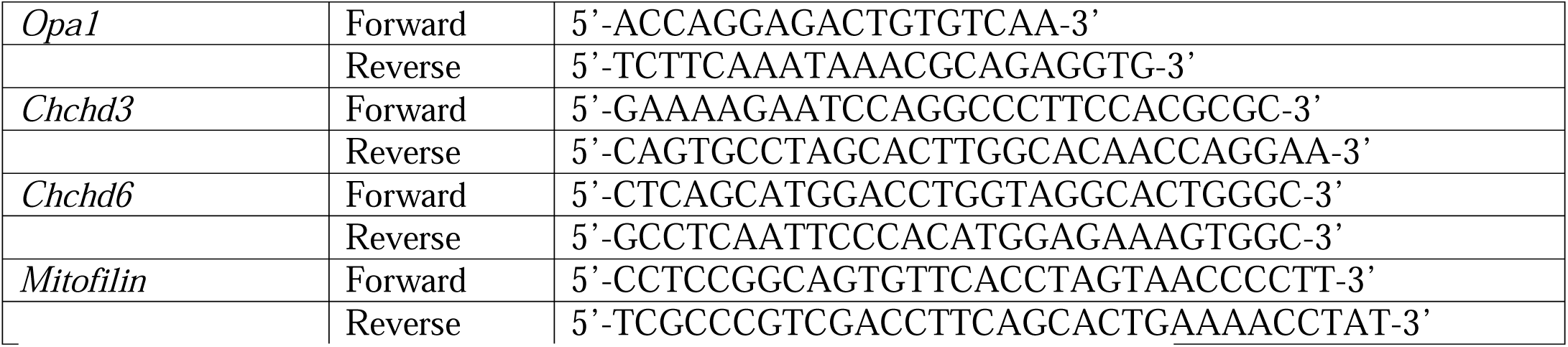
qPCR Primers Used.

**Table 2:**
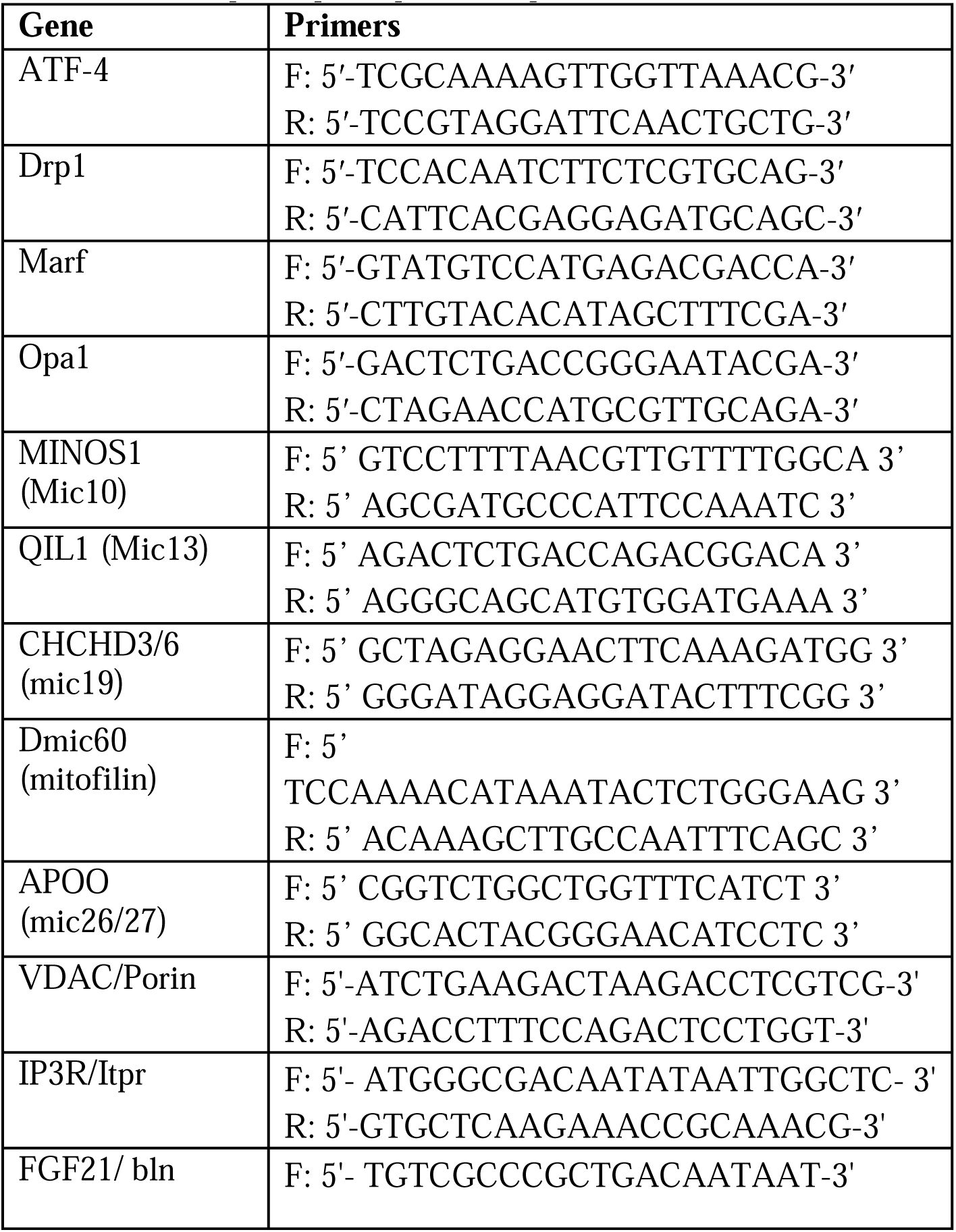
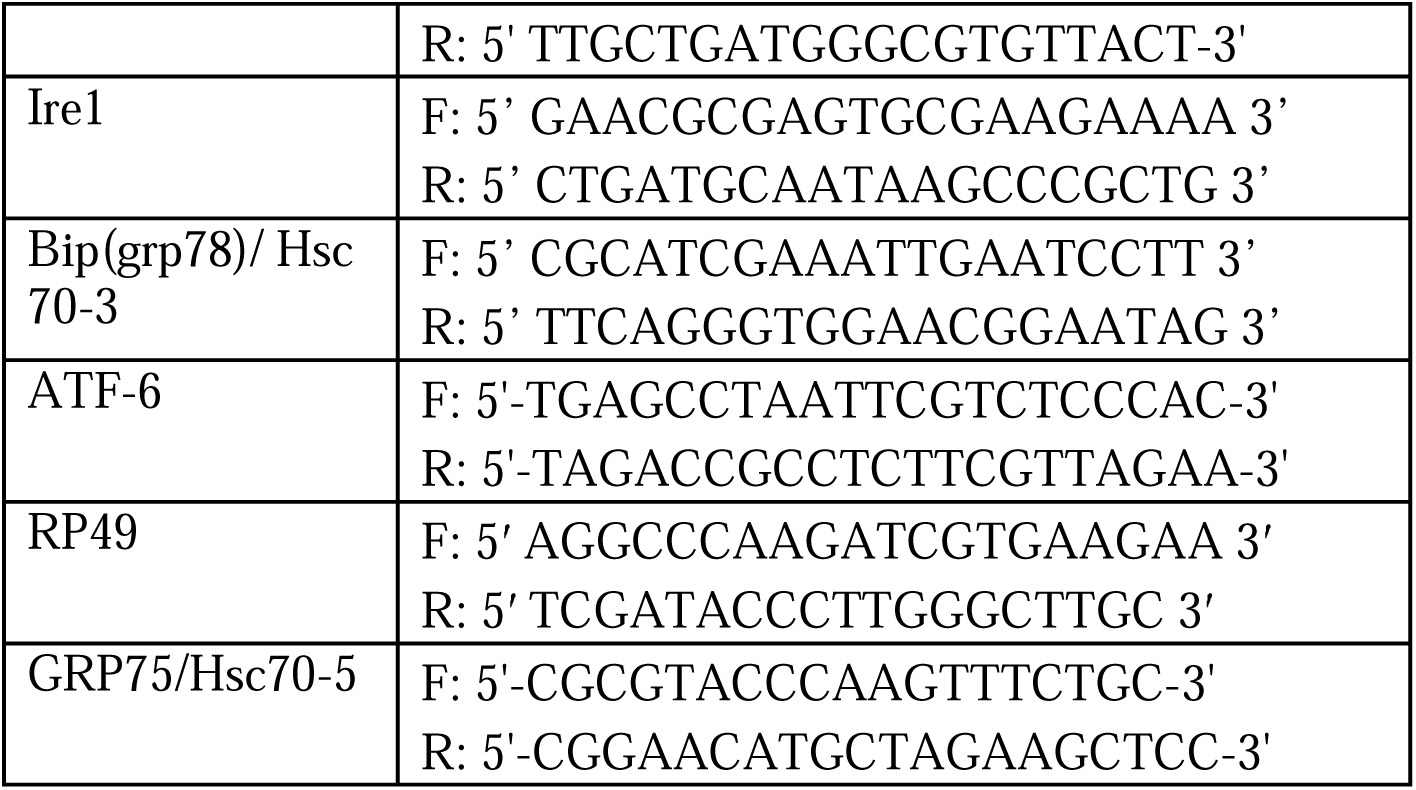
Drosophila qPCR primer sequences.

## 2D and 3D Structural Changes in Cristae and Mitochondria Exhibited Upon Loss of MICOS Complex and *Opa1*

To determine the role of OPA-1 and the MICOS complex in mitochondrial structure and networking, we ablated the genes for *Opa1* and the MICOS complex proteins in isolated primary skeletal muscle cells from 3-month-old mice. We isolated primary satellite cells, then differentiated myoblasts into myotubes. Using CRISPR/Cas9 method and a control plasmid, we knocked out the genes for MICOS complex components and *Opa1* from skeletal muscle cells.

We measured 1250 mitochondria across 10 cells, with loss of *Opa1* as a positive control for mitochondrial morphological changes, since it is well understood that *in vitro* deletion of *Opa1* alters mitochondrial morphology ^14,36,37^. Although *Opa1* expression decreases with age^38^, how loss of the MICOS complex affects mitochondria 3D morphology is poorly elucidated. Past studies have indicated knockout of MICOS subunit *Chchd3* results in fragmented mitochondria as the cristae lose their normal structure^39^. Similarly, *Chchd6* is important in maintaining crista structure and its downregulation results in hollow cristae that lack an electron-dense matrix, thereby inhibiting ATP production and cell growth^40–42^. Using transmission electron microscopy (TEM), we compared mitochondria and cristae in myotubes from wild-type (WT) and knockouts of *Opa1-*, *Mitofilin-, Chchd3-*, and *Chchd6-*knockout myotubes, which are essential for the organization of mitochondrial cristae^41,43^ (Figures 4A–E). Mitochondrial average area decreased for *Opa1-*, *Mitofilin-, Chchd3-*, and *Chchd6-*knockout myotubes (Figure 4F), while the mitochondrial circularity index (the roundness and symmetry of mitochondria) and the number of mitochondria, once normalized, increased (Figures 4G–H). This suggests that mitochondria become less complex, smaller, and more abundant upon loss of the MICOS complex. For *Opa1-*, *Chchd3-*, *Mitofilin-*, and *Chchd6-*knockouts, the number of cristae per mitochondrion decreased, as did the cristae score and cristae surface area compared with the WT (Figures 4I–K). Here cristae score is defined by the following:

**Figure 4.**
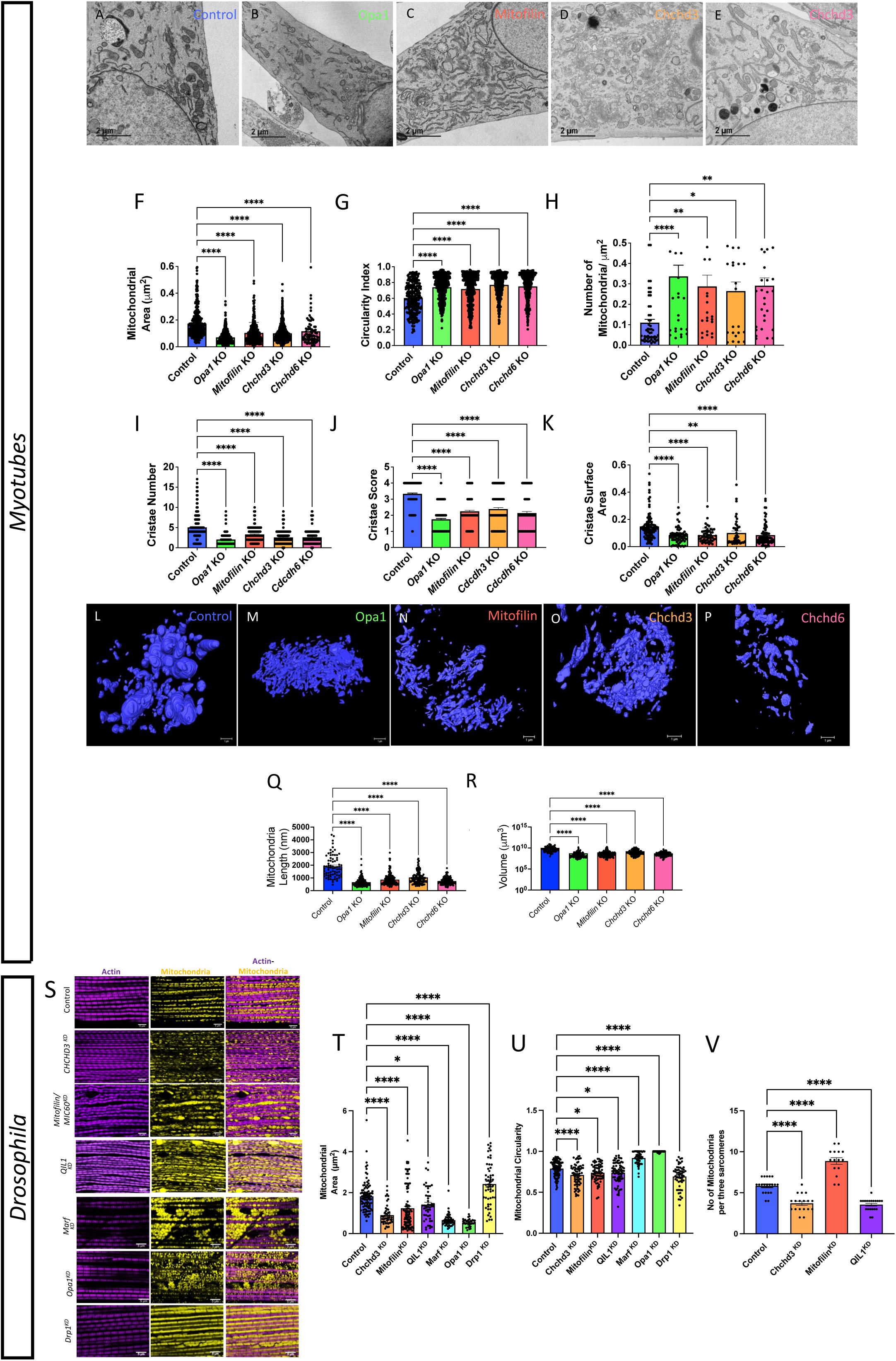
Knockout of *Opa1*, *Mitofilin, Chchd3*, or *Chchd6* in myotubes results in changes in structural changes of mitochondria and cristae in TEM and 3D reconstruction. (**A-E**) Representative images of mitochondria and cristae from myotubes of *Opa1*, *Mitofilin, Chchd3*, and *Chchd6*-knockout mice compared to WT. (**F**) Mitochondrial area in myotubes of *Opa1*, *Mitofilin, Chchd3*, and *Chchd6*-knockout mice compared to WT. (**G**) Circularity index, measuring the roundness and symmetry of mitochondria, in myotubes of *Opa1*, *Mitofilin, Chchd3*, and *Chchd6*-knockout mice compared to WT. (**H**) The number of mitochondria in myotubes of *Opa1*, *Mitofilin, Chchd3*, and *Chchd6*-knockout mice compared to WT. (**I**) Quantification of individual cristae quantity in myotubes of *Opa1*, *Mitofilin, Chchd3*, and *Chchd6*-knockout mice compared to WT. (**J**) Cristae scores measuring the uniformity and idealness of cristae in myotubes of *Opa1*, *Mitofilin, Chchd3*, and *Chchd6*-knockout mice compared to WT. (**K**) Surface area of the average cristae in myotubes of *Opa1*, *Mitofilin, Chchd3*, and *Chchd6*-knockout mice compared to WT. (**L-P**) Representative images showing 3D reconstructions of mitochondria in myotubes of *Opa1*, *Mitofilin*, *Chchd3*, and *Chchd6* knockout mice compared to WT. (**Q**) Mitochondrial 3D length in myotubes of *Opa1*, *Mitofilin*, *Chchd3*, and *Chchd6* knockout mice compared to WT. (**R**) Mitochondrial volume on log scale in myotubes of *Opa1*, *Mitofilin, Chchd3*, and *Chchd6* knockout mice compared to WT. (**S-V**) Altered *Drosophila* mitochondrial structure with loss of the MICOS complex and mitochondrial proteins. (**S**) Actin-Mitochondria staining for *Drosophila* flight tissue across knockout of MICOS complex and mitochondrial proteins. (**T**) Transmission electron microscopy (TEM) quantification of changes in *Drosophila* flight tissue the mitochondrial area, (**U**) circularity, and (**V**) quantity per sarcomere upon knockout of MICOS complex and mitochondrial proteins. Significance values * *p* ≤ 0.05, ** *p* ≤ 0.01, *** *p* ≤ 0.001, **** *p* ≤ 0.0001. Dots represent number of mitochondria quantified.

0: No sharply defined cristae are visible.

1: Over 50% of the mitochondrial area lacks cristae. 2: Over 25% of the mitochondrial area lacks cristae.

3: Irregularly shaped cristae are visible, less than 25% of mitochondria lack cristae. over 75% of the mitochondrial area.

4: Many regular-shaped cristae are visible, less than 25% of mitochondria lack cristae.

While the loss of the MICOS complex shows similar changes across all of these knockouts, in general, the least significant change was seen in the *Chchd3* knockout. Together, these data show quantitative and structural changes in both mitochondria and cristae upon loss of MICOS proteins.

TEM provides cristae detail, but not 3D morphology; therefore, we used SBF-SEM to look at the 3D structure of the mitochondria. Using myotubes with ablated genes for *Opa1* and MICOS complex subunits, as described above, we measured a total of 200 mitochondria across 10 cells. We compared mitochondria in WT, *Opa1-*, *Mitofilin-, Chchd3-*, and *Chchd6-*knockout myotubes (Figures 4L-P). We found that compared with the elongated mitochondria in the WT, the 3D length was much shorter in *Opa1* and MICOS protein knockouts (Figure 4Q). Similarly, the volume of mitochondria was also reduced in *Chchd3-*, *Chchd6-*, *Mitofilin-*, and *Opa1*- knockouts compared with the control (Figure 4R). The 3D reconstruction data, in combination with the prior TEM results, show how mitochondrial dynamics change with the loss of MICOS subunits and mimic the mitochondrial phenotypes observed across aging.

We also wanted to consider how loss of the MICOS complex and other mitochondrial genes impacted mitochondrial structure in a *Drosophila* model. Since QIL1 (Mic13) and CHCHD3/6 (mic19) were all specifically downregulated across the aging process in skeletal tissue, we sought to understand their role in *Drosophila* mitochondrial structure. We also considered how the loss of Mitofilin, which was slightly upregulated in aging, changed mitochondrial structure. Finally, we also looked at standard mitochondrial fusion proteins of OPA1 and MARF and the fission protein DRP1. Mitochondrial-actin straining showed differences in mitochondrial organization and relative Myofibrillar density (Figure 4S). From there, we utilized TEM to understand how *in vivo* flight muscle mitochondrial morphology was altered upon these KDs. As expected, loss of DRP1 caused increases in the mitochondrial area while inversely loss of MARF and OPA1 both reduced mitochondrial area (Figure 4T). Loss of all MICOS complex proteins (Mic13, Mic19, and Mic60) all showed reductions in mitochondrial volume, although this reduction was less severe in Mic13 (Figure 4T). Looking at circularity to understand mitochondrial complexity, we notice that interestingly loss of the MICOS complex paralleled the knockout of DRP1 marked by a decrease in circularity (Figure 4U). In contrast, loss of mitochondrial fusion proteins increased circularity of mitochondria. Finally, for Mic13, Mic19, and Mic60 we notably show that while the loss of Mic13 and Mic19 reduces mitochondrial quantity, yet loss of Mic60 increases mitochondrial quantity (Figure 4V). This suggests that loss of gene expressions of the MICOS complex and related mitochondrial structure may be evolutionarily conserved to some degree yet there may be certain organismal- dependent alterations in associated mitochondrial dynamics and age-related gene expression reduction.

### Changes in Oxygen Respiration Rate and Metabolite Changes Upon Loss of MICOS Complex and *Opa1*

To understand the functional impact of these structural remodeling of mitochondria upon loss of MICOS complex, we used an XF24 seahorse analyzer to measure oxygen consumption rate (OCR). Past studies have indicated that loss of *Opa1* induces bioenergetic stress and decreased electron transport chain function^37^, and ablation of the MICOS complex alters mitochondrial capacity^44,45^. We found that loss of *Opa1* or *Mitofilin* in myotubes decreased basal OCR (Figure 5A–B) and decreased ATP-linked, maximum, and reserve capacity OCR (Figure 5C–E). Although *Opa1*-knockout myotubes exhibited a decrease in proton leak, which represents protons that go from the mitochondria to the matrix without producing ATP, *Mitofilin* knockouts showed no significant difference (Figure 5F). To determine the global effects of loss of *Opa1* or the MICOS complex in skeletal muscle myotubes, we analyzed the metabolome to identify changes in metabolites that occurred with changes in mitochondria and cristae. Principal component analysis (PCA) revealed distinct populations in the control and the *Mitofilin*- knockout strains, which suggested that their genotypes contributed to the clustering (Figure 5G). To identify metabolites with the most robust ability to discriminate between true positives and false positives, we constructed a model using analysis of variance (ANOVA) to determine several key metabolites that were statistically significant (Figure 5H). This unique metabolite signature revealed that *Mitofilin* plays a critical role in regulating amino acid metabolism and steroidogenesis (Figure 5I–J). Upregulation of steroidogenesis pathways may result from the increased fluidity of membranes caused by *Mitofilin*^46,47^.

**Figure 5:**
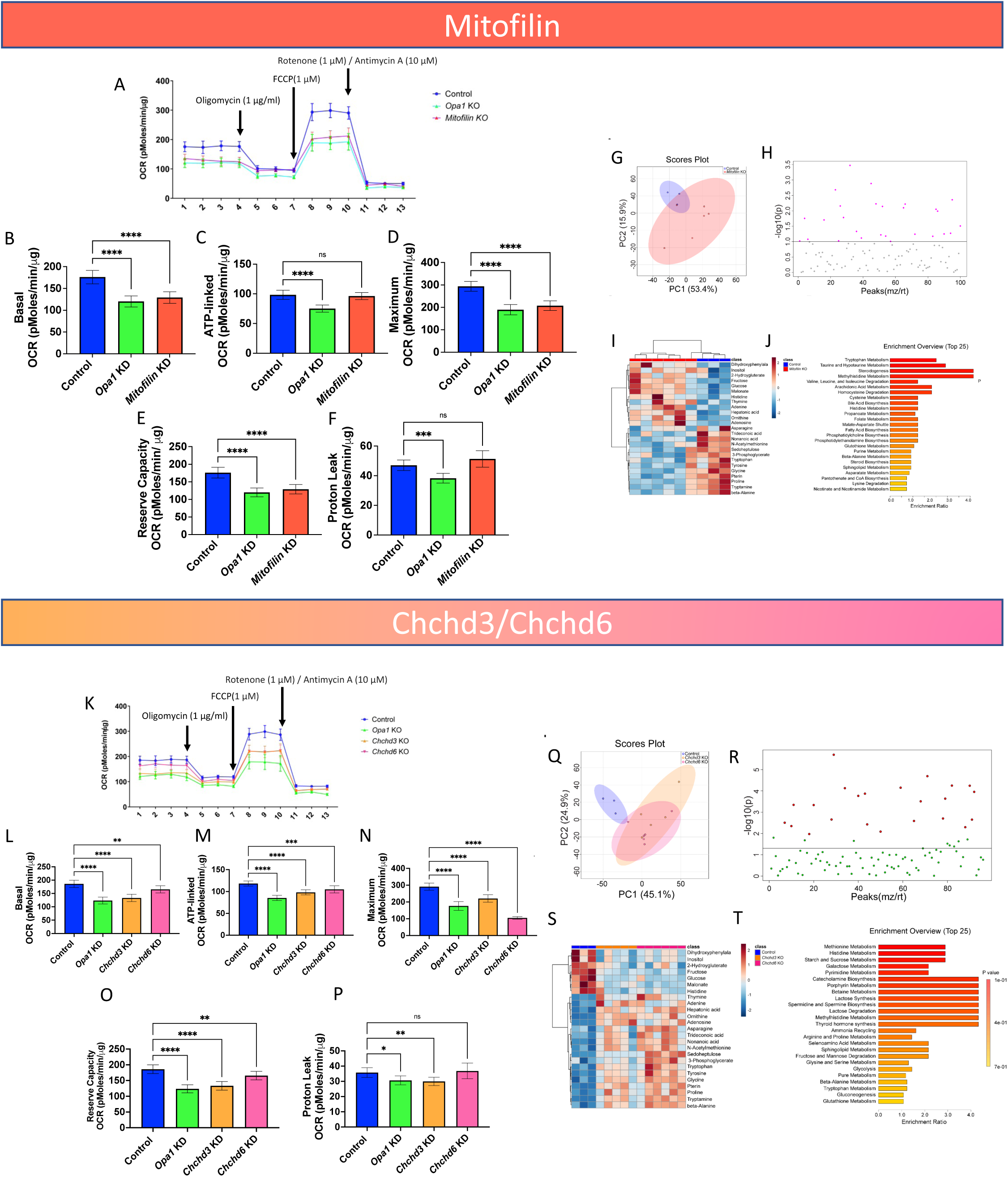
Knockout of the MICOS complex in myotubes results in changes in oxygen consumption rates and metabolomics. (**A**) OCR in myotubes of *Opa1* and *Mitofilin*-knockout mice compared to WT. (**B**) Basal OCR, the net respiration rate once non-mitochondrial respiration has been removed, in myotubes of *Opa1* and *Mitofilin*-knockout mice compared to WT. (**C**) ATP-linked respiration, shown from intervals 4−7 in the OCR graphs, was determined by the addition of oligomycin (an inhibitor of respiration), thus representing the respiration dependent on ATP, in myotubes of *Opa1* and *Mitofilin*-knockout mice compared to WT. (**D**) Maximum OCR, represented by the peak from intervals 7−11 once non-mitochondrial respiration was deducted, in myotubes of *Opa1* and *Mitofilin*-knockout mice compared to WT. (**E**) The reserve capacity, which is represented by the difference between basal OCR and maximum OCR, in myotubes of *Opa1* and *Mitofilin*- knockout mice compared to WT. (**F**) Proton leak, representing non-phosphorylating electron transfer, in myotubes of *Opa1* and *Mitofilin*-knockout mice compared to WT. (**G-J**) Metabolomic analysis upon loss of *Mitofilin* is also shown. (**G**) Metabolite PCA and (**H**) T-test comparing myotubes for control to *Mitofilin*-knockout mice. (**I**) Heatmap showing the relative abundance of ions and (**J**) enrichment analysis of metabolites, which links together several similarly functioning metabolites, with the relative abundance for *Mitofilin*-knockout. (**K**) OCR in myotubes of *Chchd3, Chchd6*, and *Opa1* knockout mice compared to WT. (**L**) Basal OCR in myotubes of *Chchd3*, *Chchd6*, and *Opa1* knockout mice compared to WT. (**M**) ATP-linked respiration in myotubes of *Chchd3*, *Cchchd6*, and *Opa1* knockout mice compared to WT. (**N**) Maximum OCR in myotubes of *Chchd3*, *Chchd6*, and *Opa1* knockout mice compared to WT. (**O**) The reserve capacity in myotubes of *Chchd3*, *Chchd6*, and *Opa1* knockout mice compared to WT. (**P**) Proton leak in myotubes of *Opa1*, *Chchd3*, and *Chchd6*, knockout mice compared to WT. (**Q-T**) Metabolomic analysis upon loss of *Chchd3* or *Chchd6* is also shown. (**Q**) Metabolite PCA and (**R**) ANOVA test comparing control to myotubes of *Chchd3* and *Chchd6* knockout mice (**S**) Heatmap showing the relative abundance of ions for control and (**T**) enrichment analysis metabolite for *Chchd3* and *Chchd6* knockout mice. Significance values \**p* ≤ 0.05, ** *p* ≤ 0.01, *** *p* ≤ 0.001, **** *p* ≤ 0.0001. For seahorse n=6 plates for experimental knockouts, while for control n=16.

In *Opa1-*, *Chchd3-*, and *Chchd6-*knockouts, there was a decrease in basal, ATP-linked, maximum, and reserve capacity OCR compared with the control (Figures 5K-O). Although proton leak OCR decreased in *Opa1-* and *Chchd3*-knockout myotubes (Figure 5P), there was no significant difference between the control and *Chchd6*. The decrease in OCR may be attributed to smaller and fragmented mitochondria; mitochondrial density decreases as fragmentation targets them for autophagy^38,48^. Together, these results show that MICOS and *Opa1* are essential for the normal respiration of muscle tissue. We also measured the effect of ablating the genes for *Chchd3* and *Chchd6* in skeletal muscle myotubes on bioenergetic metabolism. PCA revealed distinct populations in the control and the *Chchd3-* and *Chchd6*-knockouts, which showed a similar profile to that we observed in *Mitofilin* (Figure 5Q). We constructed a model using ANOVA to determine which metabolite changes in *Chchd3* and *Chchd6* knockouts were statistically significant (Figure 5R). There was a loss of protein synthesis and changes in carbohydrate metabolism (Figure 5S–T). Loss of *Opa1* typically favors fatty acid synthesis, so the results showing increased carbohydrate metabolism differ from previous *Opa1*-knockout responses^49–51^. This atypical response was evident in the increase in lactose and starch synthesis, but there was poor protein turnover, as seen in methionine metabolism (Figure 5T). Given that we saw a loss of MICOS complex proteins caused a change in metabolism, we sought to see if this change in metabolism parallels the age-related change in gastrocnemius, soleus, and cardiac metabolism.

### Metabolomics/Lipidomic Show that Across Aging, Gastrocnemius, Soleus, and Cardiac Tissue Exhibit Altered Metabolism Across Aging

Together, our results up to this point suggested an age-related change in the MICOS complex with implications for structural and functional alterations in the MICOS complex. Given we saw a loss of the MICOS complex was implicated in altered steroidogenesis and metabolism, we examined if similar pathway alterations are noted in aging to explicate future areas of study.

To investigate the factors influencing the observed changes in aged tissues, we conducted metabolomics and lipidomics analyses on both young and aged muscle tissues (Figure 6A-F). Our analysis revealed significant metabolic shifts in all three tissue types. These changes encompassed various processes, including NAD^+^ metabolism, linolenic metabolism, porphyrin synthesis, heme biosynthesis, and glycine and lysine metabolism (SFigure 3D-I). Notably, we observed an accumulation of cholic acid in aged soleus and gastrocnemius muscles (SFigure 3A, B, D, and E), a known inducer of muscle atrophy ^52^. Amino acid metabolism was dysregulated across all muscle tissues.

**Figure 6:**
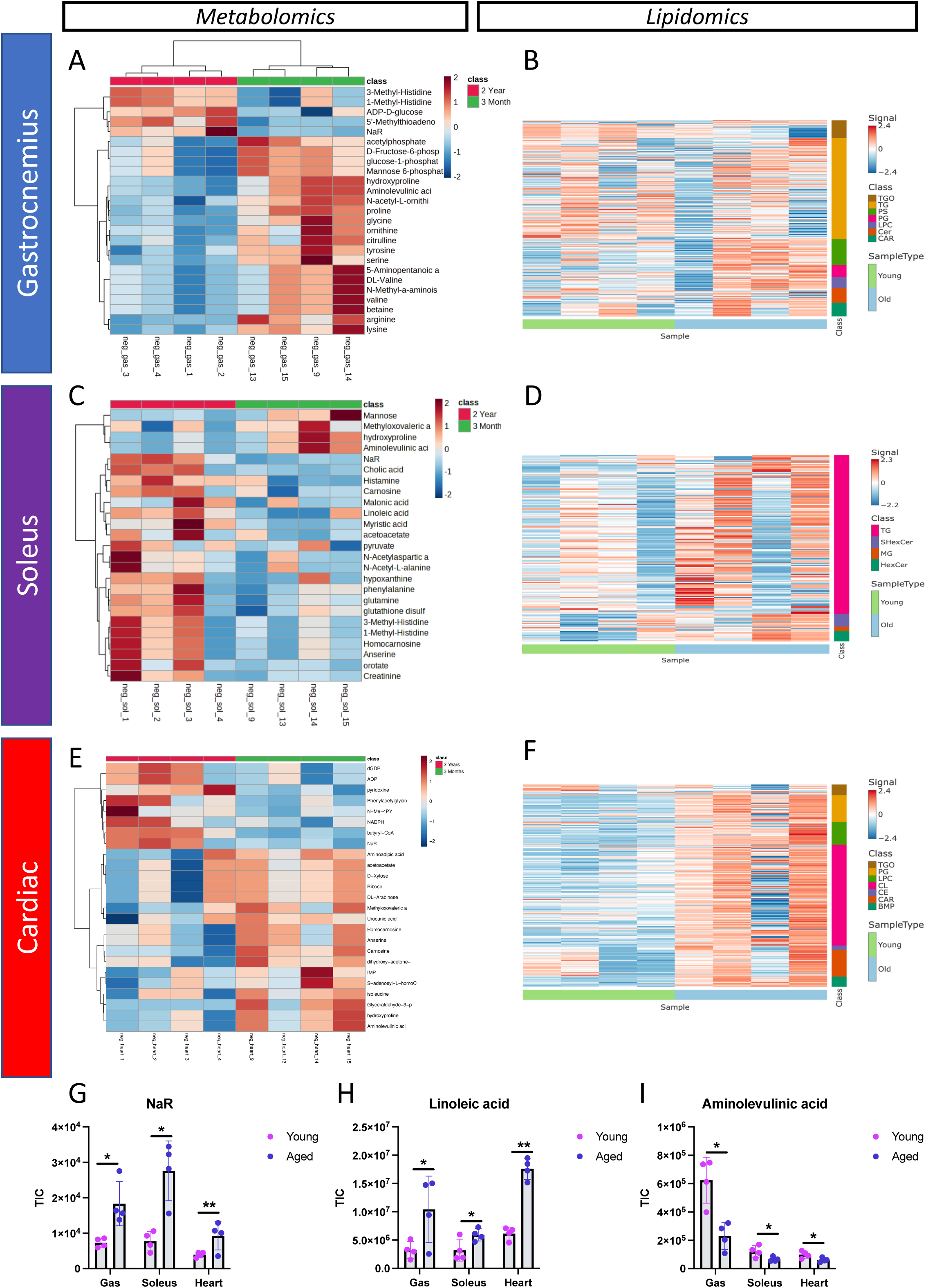
Metabolomics analysis and lipidomic profiling reveals metabolic dysregulation and disruptions in lipid classes with age in gastrocnemius, soleus, and cardiac muscles. (**A**) Metabolic heatmap representing relative abundance of metabolites and (**B**) heatmaps representing the lipidome in young and aged gastrocnemius, (**C-D**) soleus, and (**E-F**) cardiac samples. For each tissue and metabolite in the heatmaps, aged are normalized to the median of young, and then log2 transformed. Significantly differentiated lipid classes are those with adjusted p-values < 0.05 (note: p-values were adjusted to correct for multiple comparisons using and FDR procedure) and log fold changes greater than 1 or less than -1. Young, n=4; aged, n=4. For all panels, *error bars* indicate S.E.M. ***, *p* < 0.001; **, *p* < 0.01; *, *p* < 0.05; §, *p* < 0.1, calculated with Student’s *t-*test.

We conducted lipidomic profiling to investigate the role of dysregulated lipid metabolism in muscle tissues during aging (Figure 6 B, D, F; STables 1-4). Our analysis revealed notable changes in both lipid classes (SFigure 3 J-L) and lipid chain lengths (SFigure 3 M-O) with age. Among the three muscle types, the gastrocnemius and cardiac tissues exhibited the most pronounced alterations in lipid classes, while the soleus showed minimal dysregulation (SFigure 3 J-L). Additionally, the cardiac muscle displayed a higher degree of disrupted lipid chain lengths compared to the soleus and gastrocnemius muscles (SFigure 3M-O). These findings highlight the importance of altered lipid chain lengths in cellular and physiological processes, as they can affect membrane integrity, fluidity, and functionality, thereby influencing a wide range of cellular functions.

The gastrocnemius muscle exhibited dynamic changes in lipids with age (Figure 6B). These alterations affected various lipid classes, including triglycerides oligomers (TGO), triglycerides (TG), phosphatidylserine (PS), phosphatidylglycerol (PG), ceramide (Cer), and acylcarnitine (CAR). In the gastrocnemius, the levels of PS, the most abundant negatively charged phospholipid in eukaryotic membranes and enriched at cell-cell contact regions ^53^, decreased with age (SFigure 3J). Acylcarnitine, which plays a crucial role in transporting acyl groups from the cytosol into the mitochondrial matrix for beta-oxidation ^54^, was significantly reduced in aged gastrocnemius tissues (SFigure 3J). Conversely, ceramide levels increased with age in these tissues (SFigure 3J). This observation is supported by previous reports of elevated ceramide levels in replicative senescent cells, with ceramide proposed to contribute to senescence by inducing cell cycle arrest ^55^.

In the aged soleus tissues, we observed alterations in only four lipid classes (SFigure 3K). These classes included triglycerides (TG), sulfatide hexosylceramide (SHexCer), monoradylglycerol (MG), and hexosylceramide (HexCer). Notably, both HexCer and SHexCer exhibited depletion with age in the soleus (SFigure 3K). These lipids belong to the sphingolipid class, suggesting a novel role for sphingolipids in the aging process of the soleus. Additionally, MGs were found to accumulate in the aged soleus (SFigure 3K). These lipids have been linked to lipotoxicity, which triggers immunesenescence ^56^. Lastly, TGs, which are crucial for storing and transporting fatty acids within cells and in the circulation, showed a decrease in the aged soleus (SFigure 3K). This finding aligns with previous studies reporting decreased TG levels in plasma and increased levels of fatty acids during aging ^57^.

The cardiac muscle displayed age-related alterations in several lipid classes, including triglycerides oligo (TGO), phophatidyl glycerol (PG), lysophosphatidylcholines (LPC), cardiolipin (CL), cholesteryl esters (CE), acyl carnitine (CAR), and bis(monoacylglycerol)phosphate (BMP) (SFigure 3L). Notably, cardiolipins, which play a crucial role in regulating mitochondrial proteins and maintaining mitochondrial structures such as cristae and contact sites ^58^, exhibited a decrease in aged cardiac muscles (SFigure 3L). This lipid class has been implicated in age-related alterations in mitochondrial bioenergetics ^58–60^. Furthermore, we observed an accumulation of TGO in the heart, which has been linked to inflammation, endothelial dysfunction, oxidative stress, atherosclerotic plaques, and ultimately cardiovascular disease (SFigure 3L) ^61^. TGOs are closely associated with the aging process ^57^.

Returning to metabolic changes, we detected an accumulation of Nicotinic Acid Riboside (NaR) in the soleus, cardiac, and gastrocnemius tissues (Figure 6G), indicating potential compensatory mechanisms to maintain NAD^+^ levels, which are known to decline with age in muscle tissues ^62^. Furthermore, the levels of linoleic acid, an essential fatty acid crucial for cell membrane integrity and the synthesis of inflammation-related eicosanoids, increased with age in these three tissues (Figure 6H). In our metabolic profiling, aminolevulinic acid stood out as a depleted metabolite in the cardiac, soleus, and gastrocnemius tissues of aged mice (Figure 6I). Aminolevulinic acid is essential for heme production, a process vital for mitochondrial oxygen and energy production ^63^. Collectively, these profound metabolic alterations support our observations of age-related changes in these muscle tissues.

In summary, our metabolic analysis and lipid profiling unveiled significant metabolic alterations in cardiac, soleus, and gastrocnemius muscles with age, some of which may parallel the metabolic alterations seen with the loss of the MICOS complex.

## DISCUSSION

We demonstrated that either aging or loss of MICOS proteins in skeletal muscle resulted in suboptimal mitochondria morphology in a tissue-dependent manner, suggesting a potential correlation between aging and MICOS protein expression. Previous studies used 3D focused ion beam-scanning electron microscopy (FIB-SEM) to characterize the networking of the mitochondria in human^64^ and mouse skeletal muscle^65^. Another report did quantitative 3D reconstructions using SBF-SEM to define the morphological differences in the skeletal muscles of humans versus mice, and they compared patients with primary mitochondrial DNA diseases with healthy controls^25^. To the best of our knowledge, our current study is the first to use a 3D reconstruction method as an approach to studying mitochondria phenotypes in skeletal muscle aging. We prioritized using manual contour tracing as opposed to machine learning techniques to ensure the accuracy of these highly varied mitochondrial phenotypes. Future research may consider looking further at human skeletal muscle, to better compare if human gastrocnemius and soleus muscles have a similar phenotype to murine skeletal muscle. While the murine soleus is understood to have a similar transcriptome to that of humans ^66^, further elucidation of these similarities are necessary. Furthermore, considering that oxidative muscle types vary in function and structure, future studies may further seek to better discern them.

Skeletal muscle is highly mitochondrial-dependent, as well as mitochondria enriched, which comprise ∼6% of the cell volume and changes with aging^67^. Gastrocnemius muscle has both type I slow-twitch muscle fibers and type II fast-twitch muscle fibers; type I fibers are more effective for endurance, while type II fibers better support short bursts of muscle activity^21,67,68^. Of relevance, sarcopenia is characterized by both types reducing in frequency and size while type II fibers transition to type I ^69^. In contrast, other forms of muscle degeneration such as disuse atrophy does not see a change in fiber number and an inverse shift from type I fibers to type II ^69^. This suggests it is possible that age-related changes in mitochondrial shape may be due to sarcopenia-dependent alterations in fiber frequency, which may have different mitochondrial phenotypes. However, our 3D morphologic data did not permit discrimination of the two fiber types, although we observed many variable muscle fibers within a sample and between all differently aged cohorts. Past studies in human skeletal muscle separated these by taking two portions of samples and assessing fiber type via SDS-PAGE^70^. Importantly, past studies have found rather large differences in mitochondrial content and morphology in chicken muscle fiber subtypes^71^. In a chicken model, past literature has found that type I does not contain lipid droplets, and used the presence of lipid droplets to identify type II fibers ^71,72^. However, to our knowledge, this technique has not yet been applied to the 3D reconstruction of aged murine skeletal muscle models.

Using 3D reconstructions, we found that all parameters of the mitochondria changed during aging, including smaller volume, area, and perimeter (Figure 1A-X), and the mitochondria became less interconnected in gastrocnemius while other tissue types saw a less significant decrease in volume. This increased fragmentation suggests a decreased mitochondrial fusion, in association with age-dependent decreases of OPA-1, which is a regulator of mitochondrial fusion. We also saw a decrease in the MCI during aging, suggesting that mitochondria decrease in networking, but their shape may not radically change as they increase in sphericity as they age (Figure 2).

MICOS proteins play key regulatory roles in mitochondrial structure and function^17,27,48^. We determined by TEM 3D reconstructions that the loss of *Mitofilin*, *Chchd3*, and *Chchd6* resulted in fragmentation, disrupted cristae, and smaller mitochondria (Figure 4), similar to the loss of *Opa1*, which is known to cause changes in oxidative phosphorylation ^17,30,37^. Overall, mitochondria lacking the MICOS genes had characteristics similar to those of aged mouse skeletal muscle (Figures 1−2). This similarity in phenotype suggests there may be an association. Thus, changes in mitochondrial morphology due to aging may be caused by a lack of MICOS protein expression. This is supported by the RT-qPCR data that showed decreased *Chchd3*, *Chchd6*, *Mitofilin*, and *Opa1* transcription with aging (Figure 3). Notably, however, while both tissue types exhibit diminished mRNA transcripts, only skeletal muscle shows diminished protein levels of the MICOS complex and OPA1 (Figure 3). This may be a cell-specific response, exhibiting down-regulated proteins in cardiomyocytes but more abundantly expressed proteins in fibroblasts ^73^. This may also represent a compensatory mechanism, where the half-life of the protein is much longer in cardiac tissue to buffer against increased loss of mRNA transcripts. We also note that there are differential metabolomics and lipidomics across different tissue types, which suggests that this MICOS-complex-dependent response to aging may differ across cell types due to altered metabolomics. In any case, the tissue-specific loss of the MICOS complex must be further elucidated.

In *Drosophila*, since *Mitofilin* is slightly upregulated (Figure 3L), its expression may act as a counteractive mechanism relative to the loss of other MICOS complex proteins across the aging process. Notably, in flies, we also show that *Mitofilin* loss increases mitochondrial quantity conversely to the loss of other MICOS complex components. This suggests that the roles of *Mitofilin* may be differential in Drosophila and murine models, highlighting the need for additional studies on *Mitofilin*. Furthermore, the differential roles of the MICOS complex proteins need further elucidation. While there are commonalities in phenotypes between all MICOS complex components when knocked down, we see that loss of *Mitofilin* and *Chchd6* do not have the same reduction in proton leak as *Chchd3* KD (Figure 5F, P). While past studies have indicated that all three of these components are necessary for the stabilization of the MICOS and SAM complex, and consequently cristae architecture ^27,39–41,74^, our findings suggest greater research is necessary to understand if compensation for specific components of the complex avoids dysfunctional phenotypes upon loss of certain MICOS complex components.

Although there is a link between aging and loss of *Opa1* ^31,38^, little is known about the role of the MICOS complex in aging. Changes in mitochondrial architecture and loss of integrity may be caused by decreased MICOS proteins; thus, it will be important to develop methods to restore MICOS proteins and *Opa1* lost during aging to decrease the deleterious effects of mitochondrial dysfunction. Although studies have examined the role of knockouts on mitochondrial dynamics, few studies have examined how the loss of MICOS proteins may be restored to mitochondria^30,75^. To better elucidate potential future studies, we sought to consider the similarities in metabolomics and lipidomics in loss of the MICOS complex and the aging process.

We noticed differential changes in metabolomics and lipids both in comparison between skeletal and cardiac tissue, but even between soleus and gastrocnemius (Figure 6). This highlights the importance of considering tissue-dependent differences, which may arise due to the tissue-dependent mitochondrial structures we observed. Metabolomics showed that several pathways acted to cause muscular dystrophy while other pathways such as Nicotinic Acid Riboside acted to rescue these pathways. Cardiac tissue shows the largest change in lipid alterations across aging, which suggests that lipid accumulation may play a larger role in cardiac, which matches prior literature which shows that lipid metabolism is closely linked with pathology ^76^. Conversely, soleus shows a Sphingolipid decrease, which may show that the soleus age due to lipo-toxicity. Sphingolipid also is associated with MERC regions during and prior to apoptosis ^77^. This highlights the importance of future studies examining mitochondria-lipid droplet associations, or interactions with ER ^78^ to see if these contact sites change across the aging process to protect against lipotoxicity, as well as experiments employing mass spectrometry imaging to better understand the spatial distribution of these metabolites across aging ^79^. In gastrocnemius tissue, we observe numerous phospholipids decrease (Figure 6). We also saw a loss of linoleic acid, which is important for membrane integrity ^80^. This suggests altered mitochondrial membrane viscosity across the aging process. Past findings have also observed age-dependent increases in viscosity which affect oxidative phosphorylation through modulation of supercomplexes performance ^81^. Notably, decreased membrane viscosity occurs in age-related pathologies including Alzheimer’s Disease, highlighting that these changes may have implications in pathomechanisms ^82^. One potential target is the MICOS complex, as past studies highlight that lipids, including cardiolipin, can interact with cristae ^83^. While we only see a change in cardiolipins in cardiac tissue, it is possible that since this region has the least drastic change in mitochondrial structure, cardiolipins protect against MICOS-dependent loss in structure. It is possible that when you lose cristae morphology, you may increase in cardiolipins. In conclusion, we can conclude that aging may also be due to alterations in membranes mediated by lipids, affecting the MICOS complex and modulating mitochondrial structure and function.

Some of these age-related lipidomic and metabolomic changes may be due age-dependent alterations in the MICOS complex. Many studies have analyzed the mitochondrial metabolome using mouse skeletal muscles^67,84–88^. We found that loss of *Mitofilin* affected cristae morphology (Figures 4I–K), decreased oxidative phosphorylation (Figure 5A), and may have increased lipid and steroid synthesis, which may be important for MERCs regulation and cristae formation. We found an increase in tryptophan and methyl histidine metabolism (Figure 5J) and an increase in taurine metabolism and hypotaurine, a key sulfur-containing amino acid for fat metabolism. Loss of *Opa1* also changes amino acid and lipid metabolism, similar to the loss of *Mitofilin*^49–51^. Steroidogenesis, which makes the membrane less rigid, was increased. Since the loss of *Mitofilin*, *Chchd6*, or *Chchd3* caused a loss of oxidative capacity (Figure 5A-F; 5K-P), increased steroid synthesis may allow the cell to recover bioenergetic functions, as steroid compounds decrease membrane viscosity, with the incorporation of estrogen^46^. In the absence of *Mitofilin*, cristae junctions and contact sites fall apart^89^; thus, *Mitofilin* is critical for maintaining cristae^90,91^. Cells lacking *Mitofilin* may make steroids to help the membrane to reconfigure broken cristae. Since the loss of *Opa1* causes more MERCs^92^, loss of *Mitofilin* may increase phospholipids (Figure 5J) because of increased smooth MERCs, which are associated with lipid changes^93^. This is supported by the fact that biosynthesis of phosphatidylethanolamine and phosphatidylcholine, and metabolism of arachidonic acid and sphingolipid, increased with loss of *Mitofilin* (Figure 5J). Since these phospholipids aggregate around MERCs and may shuffle into the ER, *Mitofilin* may be a critical gene for regulating cristae morphology, with a key, novel role in regulating mitochondrial metabolism.

*Mitofilin* may be an important target in the future to restore energy production. Loss of *Mitofilin* may lead to ER stress, which, via ATF4, activates amino acid transporters^94^ that further activate mTORC1. ER stress activates mTORC as a result of a decrease in glucose^88^. Critically, mTORC1 affects glucose homeostasis^95^, which may lead to inefficient energy use. This can result in changes in autophagy. Therefore, as a downstream effect of *Mitofilin* loss increasing mTORC1, this may explain why deletion of MICOS in *Drosophila* increases autophagy^48^.

Similarly, loss of *Opa1* increases ER stress^37^, and loss of *Mitofilin* may cause a similar increase in amino acid catabolism. If ER stress activates amino acid transporters, branched-chain amino acids could increase ER stress, resulting in a positive feedback loop that affects the health of the cell, cellular energy, metabolism, and antioxidants. ER stress may also be responsible for the poor performance and fragmentation of mitochondria (Figure 4-5). Loss of *Mitofilin* may result in the breakdown of protein pathways that regulate ER stress. Other amino acid pathways, such as homocysteine (Figure 5J), are involved in triglyceride uptake and increased intracellular cholesterol, suggesting that proteins like ATF4^88^ and the MICOS complex^13,27^ are important for aging. In particular, the MICOS components may prevent mitochondrial fragmentation by blocking ER stress pathways in aging. We notably show that several MERC proteins are differentially regulated concomitantly with MICOS complex proteins across the aging process in *Drosophila* (Tables 1-2). Further studies are needed to better understand the role of MICOS in MERC formation and the relationship between smooth MERC and lipid synthesis.

Although *Mitofilin* is a key component of the MICOS complex, other components are likely also important. The loss of *Chchd3* or *Chchd6* leads to a decrease in and disassembly of all *Mitofilin* subcomplex components in mammals, with abnormal cristae morphology and growth defects^28,29,40,74,96–98^. Downregulation of *Chchd3* is linked to type 2 diabetes^99^. In our metabolomics enrichment dataset (Figure 6T), loss of *Chchd3* or *Chchd6* in mouse myotubes resulted in a preference for alternative energy sources, such as lactate, lactose, and starches.

Supplementation of healthy myotubes with galactose leads to a 30% increase in oxidative capacity (i.e., OCR) due to an increase in AMPK phosphorylation and cytochrome c oxidase (COX) activity, thereby forcing cells to become more oxidative to maintain ATP levels^100^. In our samples, as oxidative metabolism decreased, anaerobic metabolism and lactate levels increased, forcing cells to produce ATP by anaerobic glycolysis. However, long and high-level exposure to D-galactose generates free radicals, which alter MERCs, causing mitochondrial dysfunction and inducing aging^101,102^. This is the likely explanation for mitochondrial fragmentation in aged samples and loss of the MICOS complex, which should be investigated further.

In conclusion, we present a quantitative evaluation of mitochondrial morphology in mouse skeletal muscle and cardiac tissue using 3D reconstructions, while TEM was utilized in cell lines to understand factors including cristae architecture. We found structural changes, including in gross 3D mitochondrial structure, which conferred functional differences upon loss of MICOS proteins. Similar changes in mitochondrial morphology were observed in aging muscles and for loss of MICOS proteins in mouse skeletal muscle, and MICOS proteins decreased with age. We also found that metabolomics and lipidomics are heavily altered across the aging process, pointing towards the potential roles of the MICOS complex in membrane integrity across the aging process which must be further investigated. *In vivo* Drosophila models also highlight the importance of understanding the tissue-dependent response to aging, roles of individual components in the MICOS complex, and potential MICOS complex-MERC pathway interactions that may regulate mitochondria and structure. This suggests a relationship between the MICOS complex and aging, and further studies using 3D reconstruction could elucidate the link between sarcopenia, the MICOS complex, and disease states in mitochondria.

## EXPERIMENTAL PROCEDURES

### Animal Care and Maintenance

All procedures for the care of mice were in accordance with humane and ethical protocols approved by the University of Iowa Animal Care and Use Committee (IACUC) following the National Institute of Health (NIH) Guide for the Care and Use of Laboratory Animals as described previously^37^. Therefore, all studies are performed in accordance with the ethical standards laid down in the 1964 Declaration of Helsinki and its later amendments. All experiments used WT male C57Bl/6J mice housed at 22 °C on a 12-hour light, 12-hour dark cycle with free access to water and standard chow. Mice were anesthetized with 5% isoflurane/95% oxygen.

### RNA Extraction and RT-qPCR

Total RNA was extracted from tissue using TRIzol reagent (Invitrogen, cat #), purified with the RNeasy kit (Qiagen Inc, cat #), and quantitated by the absorbance at 260 nm and 280 nm using a NanoDrop 1000 (NanoDrop products, Wilmington, DE, USA) spectrophotometer. Total RNA (∼1 µg) was reverse transcribed using a High-Capacity cDNA Reverse Transcription Kit, (Applied Biosciences, Carlsbad CA, cat #) followed by real-time quantitative PCR (qPCR) reactions using SYBR Green (Life Technologies, Carlsbad, CA, cat #)^103^. Triplicate samples for qPCR (∼50 ng) in a 384-well plate were placed into ABI Prism 7900HT instrument (Applied Biosystems) programmed as follows: 1 cycle at 95°C for 10 min; 40 cycles of 95°C for 15 s; 59°C for 15 s, 72°C for 30 s, and 78°C for 10 s; 1 cycle of 95°C for 15 s; 1 cycle of 60°C for 15 s; and one cycle of 95°C for 15 s. Data were normalized to glyceraldehyde-3-phosphate dehydrogenase (*Gapdh*), and results are shown as fold changes. qPCR primers were designed using Primer-Blast or were previously published sequences^37^ as shown in Table 1.

### Experimentally evolved Drosophila populations, RNA extraction, qPCR

For this work, we used groups of experimentally *Drosophila melanogaster* where hundreds of generations of selection on reproductive timing has produced populations with markedly different aging an longevity patterns. In the control populations, termed CO_1-5_, genetically diverse populations (census size ∼2000 per replicate) are maintained on a 28-day generation cycle. From these CO populations, a set of population maintained on a 9-day cycle were created – ACO_1-5_ (See ^104^ for more details on how populations were created and still maintained). As a result of this difference and hundreds of generations, the ACO populations have evolved to optimize for early reproduction and now develop more quickly and die at much younger ages than their controls ^34^. We also know that the accelerated aging seen in the ACO flies is underlain by significant genetic differentiation ^105^, differences in patterns of gene expression ^35^, and metabolomic differentiation ^33^. The ACO populations are also referred to in this paper as “aged flies”, whereas the CO populations are referred to as “control flies”. As such, we believe comparisons between them can yield insights into the factor that broadly shape differences in rates of senescence between individuals.

Here we use qPCR to compare gene expression for the following genes (Table 2). Specifically, we looked at differences in cardiac tissue from 21-day old flies from the ACO and CO populations. We chose day 21 based on demographic data ^34^ and whole-body transcriptomic data^35^ showing that cohorts are especially differentiated at this age. For each group, we collected heart tissue using the following protocols. On day 21 from egg, female fruit flies from each of the 10 ACO and CO populations were anesthetized using Fly Nap (Carolina, NC, USA), a triethylamine-based anesthetic, for about 1-minute or until no movement was detected. Flies were dissected in oxygenated artificial hemolymph to expose the cardiac tubes. Fly abdomens were cut open in order to remove guts/intestines, fat, and ovaries. After removal, hearts were exposed and additional fat and pericardial cells were carefully suctioned away from the cardiac tube ^106^. It is not possible to fully remove all fat and pericardial cells from the cardiac tube without damaging the cardiac tube itself. Approximately 18-20 hearts from adults were pooled for each biological replicate. Three biological replicates were collected for each ACO and CO population. Total RNA was extracted using Qiazol (Qiagen) and miRNeasy Mini Kit (Qiagen, catalog no. 217004). An optional on-column DNase digestion was also performed since we expected our samples to contain less than 1 ug total RNA (Qiagen RNase-Free DNase set, cataolog no. 79254). RNA was reverse transcribed to cDNA using a QuantiTect Reverse Transcription Kit (Qiagen).

All cDNA samples were diluted to a standard concentration of 1ng/µl prior to data collection. Three replicates from each population (ACO and CO) were arbitrarily selected after filtering for DNA concentration and purity (260/280 ratio >1.8). Samples were run in duplicate on a CFX96 Touch platform (Bio Rad Laboratories, Hercules, CA). Each well contained 7.5 uL of 2X iTaq SYBR Green Supermix (Bio Rad Laboratories, Hercules, CA), 3 uL of ddH2O, 0.75 uL of each primer, and 3 uL of template cDNA. The protocol for each primer pair was as follows: 1 cycle at 95°C for 2 minutes, 40 cycles of 95°C for 30 seconds, 60°C for 30 seconds after which a melt curve analysis was conducted for quality control. Genes dMic60, Bip, and Ire1 were run at an annealing temperature of 62°C. Data was analyzed using the ΔΔCt method and normalized to the *Drosophila* gene rp49/RpL32. Primers were designed using Primer Blast or previously published sequences as shown in Table 3.

**Table 3:**
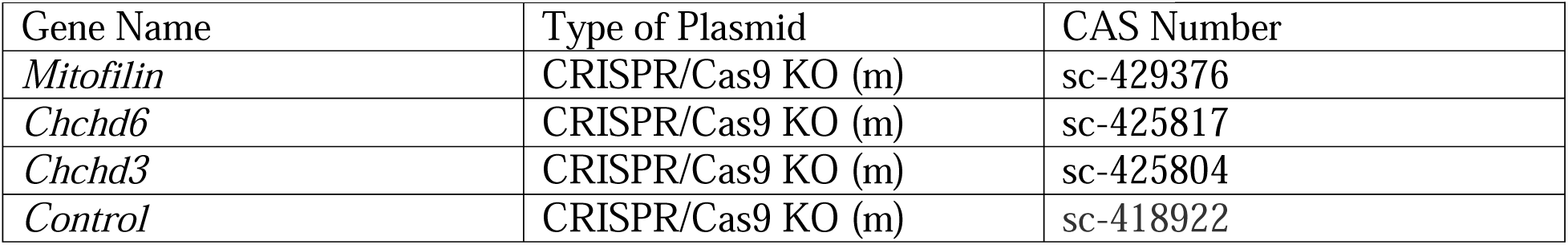
Guide RNA and Plasmids Used.

### Drosophila strains and genetics

Genetic crosses were conducted using yeast corn medium at a consistent temperature of 25 °C, unless otherwise specified. The Mef2-Gal4 was employed as a control within their respective genetic backgrounds. Both male and female subjects were used and analyzed collectively, as no discernible sex differences in mitochondrial morphology were observed in wild-type muscles. Mef2-Gal4 (III) was utilized for muscle-specific knockdown of MICOS genes. The mitochondrial network was visualized using UAS-mito-GFP, located on the second chromosome (BS# 8442).To achieve muscle-specific knockdown MICOS genes, UAS-RNAi trip lines were used for UAS- *Chchd3* RNAi (BS#51157), UAS-*Mitofilin* RNAi (BS# 63994), UAS-QIL1 RNAi (BS# 44634), UAS-Drp1 RNAi (BS# 51483), UAS-Marf RNAi (BS# 55189), and UAS-Opa1 RNAi (BS#32358). All stocks were acquired from the Bloomington Drosophila stock center, denoted by the Bloomington Stock Number (BS#). All chromosome designations and gene symbols are mentioned in FlyBase ((http://flybase.org)).

### Mitochondrial staining

Adult Drosophila thoraces, aged 2-3 days, were dissected in 4% paraformaldehyde (PF, Sigma) using fine scissors and processed as mentioned in Katti et al^1^. The dissection process isolated including Indirect Flight Muscles (IFMs). Muscles were then fixed in 4% PF for a specific duration: IFMs for 1.5 hours. This fixation process was performed using a rotor, followed by a washing sequence in PBSTx (PBSLJ+LJ0.3% TritonX100) thrice, each time for 15 minutes. Mitochondria were visualized using mitochondrial staining dyes, either Mito Tracker Red (M22425, Thermofisher, USA) or DMef2-Gal4 driven UAS-mito-GFP. Actin staining was conducted by incubating the muscles in 2.5 µg/ml of Phalloidin in PBS (Sigma, 1 mg/ml stock of Phalloidin TRITC) at 25 °C. there incubated muscles for 40 minutes at room temperature. Following staining, tissues were mounted on a glass slide using Prolong Glass Antifade Mountant with NucBlue stain (P36985, Thermofisher, USA). Images were subsequently captured using a Zeiss 780 confocal microscope.

### Mitochondrial measurements

Measurements were carried out following procedures previously established^1,2^. Images were loaded into the ImageJ software (https://imagej.net), a freely available image processing program. Individual mitochondria were outlined using the freehand tool on 2D light microscopic images provided by the software. The area and aspect ratio (the ratio of the major axis to the minor axis) of each mitochondrion were then calculated using the same ImageJ software. For each data set, three animals (n=3 animals) were analyzed, and these analyses were part of three independent experiments conducted to gather quantifiable data. For further analysis, the number of mitochondria was counted across every three-sarcomere segment.

### Isolation of Satellite Cells

Satellite cell differentiation was performed as previously described^14,37^. Gastrocnemius muscles were dissected from 8−10 week-old WT mice and washed twice with 1× PBS supplemented with 1% penicillin-streptomycin and Fungizone (300 mL/100 mL). DMEM- F12 medium with collagenase II (2 mg/mL), 1% penicillin-streptomycin, and Fungizone (300 mL/100 mL) was added to the muscle which was agitated for 90 min at 37°C. For the second wash, collagenase II was changed to 0.5 mg/mL, and the muscle was agitated for 30 min at 37°C. The tissue was cut, passed through a 70 mm cell strainer, and after centrifugation satellite cells were plated on BD Matrigel-coated dishes. To differentiate cells into myoblasts, a mixture of DMEM-F12, 20% fetal bovine serum (FBS), 40 ng/mL basic fibroblast growth factor (bFGF, R and D Systems, 233-FB/CF), 1× non-essential amino acids, 0.14 mM β-mercaptoethanol, and 1× penicillin/streptomycin, and Fungizone was used. Myoblasts were maintained with 10 ng/mL bFGF and when cells reached 80% confluence, myoblasts were differentiated in DMEM-F12, 2% FBS, 1× insulin–transferrin–selenium medium. Cells were cultured at 37°C, 5% CO_2_ Dulbecco’s modified Eagle’s medium (DMEM; GIBCO) supplemented with 10% FBS (Atlanta Bio selected), and 1% penicillin-streptomycin (Gibco, Waltham, MA, USA).

### CRISPR-Cas9 Knockouts

After three days, myotubes were incubated with CRISPR/Cas9 mixtures to produce the following knockouts—control CRISPR/Cas9 (sc-418922), *Chchd6* (Mic25) CRISPR (sc- 425817), *Chchd3* (Mic19) CRISPR (sc-425804), and *Mitofilin* (Mic60) CRISPR (sc-429376) (Santa Cruz Biotechnology, California, US), with the use of guide RNA (Table 2). We incubated 2.5% relevant CRISPR, 2.5% RNAiMax (ThermoFisher Scientific; cat # 13778075), and 95% Opti-MEM (Gibco; cat #31985070) in a tube for 20 minutes. Cells were washed twice with PBS after removal of the medium, then 800 μL of OPT-MEM and 200 µL of the CRISPR mixture were added to each well and ran in triplicates. Cells were incubated for 4 hours at 37 C, 1.0 mL of DMEM medium was added, cells were incubated overnight. The myotubes were then washed with PBS and the medium was replaced. Experiments were performed between 3 and 7 days after knockout for a total of 6 days of differentiation.

### Serial Block-Face Scanning Electron Microscope (SBF-SEM) Processing of Mouse Muscle Fibers

SBF-SEM preparation was performed as described previously^15,36,107^. Male mice were anesthetized with 5% isoflurane. Once the hair and skin were removed, the hindlimbs were incubated in 2% glutaraldehyde with 100 mM phosphate buffer for 30 min. Gastrocnemius muscles were dissected, cut into 1-mm^3^ cubes, and incubated in 2.5% glutaraldehyde, 1% paraformaldehyde, 120 mM sodium cacodylate solution for 1 hour. Tissues were washed three times with 100 mM cacodylate buffer at room temperature before immersion in 3% potassium ferrocyanide and 2% osmium tetroxide for 1 hour at 4°C, then treated with 0.1% thiocarbohydrazide, 2% filtered osmium tetroxide for 30 min, and left overnight in 1% uranyl acetate at 4°C. Between each step, three de-ionized water washes were performed. The following day, samples were immersed in 0.6% lead aspartate solution for 30 min at 60°C and dehydrated in graded concentrations of acetone. Dehydration was for five min each in 20%, 50%, 70%, 90%, 95%, and 100% acetone. Tissues were impregnated in Epoxy Taab 812 hard resin, then embedded in fresh resin, and polymerized at 60°C for 36–48 hours. Once polymerization was complete, blocks were sectioned for TEM to identify areas of interest, trimmed to 0.5 mm × 0.5 mm, and glued to aluminum pins. Afterward, pins were placed in an FEI/Thermo Scientific Volumescope 2 SEM, a state-of-the-art SBF imaging system. Running on a FEI/Thermo Scientific Volumescope 2 SEM, a state-of-the-art SBF imaging system, we obtained 300−400 10 µm by 10 µm ultrathin (90 nm) serial sections per previous techniques ^15^. All sections were collected onto formvar-coated slot grids (Pella, Redding CA), stained, and imaged as previously described^15,36,107^.

### Quantification of TEM Micrographs and Parameters Using ImageJ

Quantification of TEM images was performed as described previously using the NIH *ImageJ* software^14,36^. Cells were divided into four quadrants and two quadrants were selected randomly for complete analysis. Individuals blinded to the experimental design had a minimum of 10 cells to measure with three analyses to obtain accurate and reproducible values. If variability occurred, the number of cells was expanded to 30 cells per individual to reduce the variability.

### Measurement of OCR Using Seahorse

To measure cellular respiration, the XF24 extracellular flux (XF) bioanalyzer (Agilent Technologies/Seahorse Bioscience, North Billerica, MA, USA) was used. Cells were plated at a density of 2 × 10^4^ per well and differentiated. After 3 days of differentiation, *Opa-1*, *CHCHD3*, *CHCHD6*, or *Mitofilin* genes were knocked out as described above. Three days after knockdown, the medium was changed to XF-DMEM, and cells were kept in a non-CO_2_ incubator for 60 min. The basal OCR was measured in XF-DMEM. Oxygen consumption was measured after the addition of oligomycin (1 μg/mL); carbonyl cyanide 4-(trifluoromethoxy)phenylhydrazone (FCCP; 1 μM); rotenone (1 μM) and antimycin A (10 μM) ^37,108^. Cells were then switched to glucose-free XF-DMEM and kept in a non-CO_2_ incubator for 60 min for the glycolysis stress test. Seahorse experimental data are for triplicate Seahorse plates. Three independent experiments were performed with four to six replicates for each time and for each condition and representative data from the replicates are shown.

### Segmentation and Quantification of 3D SBF-SEM Images Using Amira

Intermyofibrillar (IMF) mitochondria are located between myofibrils, arranged in pairs at the z-band of each sarcomere, with 2D elongated tubular shapes^109^, and these were the mitochondria analyzed. For each region of interest across the three age groups, we analyzed 300 slices at 50 µm intervals in transverse intervals. For 3D reconstruction, SBF-SEM images were manually segmented in Amira as described previously^15,36^. All serial sections (300−400 slices) were loaded onto Amira and structural features were traced manually on sequential slices of micrograph blocks. Structures in mice were collected from 30–50 serial sections that were then stacked, aligned, and visualized using Amira to make videos and quantify volumetric structures. An average of 500 total mitochondria across four ROIs from three mice were collected for quantification. For 3D reconstruction of myotubes, approximately 20 mitochondria from a minimum of 10 cells were collected. Quantification of SBF-SEM images was performed as described previously^15^ using the Amira software (Thermo Scientific).

### Western-Blotting

Tissues from adult (3 months) and aged (2 years) mice were lysed with RIPA lysis buffer (1% NP40, 150 mM NaCl, 25 mM Tris base, 0.5% sodium deoxycholate, 0.1% SDS, 1% phosphatase inhibitor cocktails #2 (Sigma P5726-1ML) and #3 (Sigma P0044-1ML), 1 cOmplete protease inhibitor tablet (Sigma 04693159001)). Protein content was quantified using a BCA Assay (Thermo Scientific VLBL00GD2) and equal protein was run on 4-20% Tris-Glycine Gels (Invitrogen WXP42012BOX). Protein was transferred to a nitrocellulose membrane (Li-Cor 926- 31092). Membranes were incubated overnight with primary antibodies overnight at 4 °C: MTOC1 (Invitrogen PA5-26688), phospho S406 ATGL (Abcam ab135093), DRP1 (CST 8570S), pDRP1 (CST 6319S), OPA1 (BD Biosciences 612306), Mic60/mitofilin (Abcam ab110329), SLC25A46 (Abcam ab237760), SAM50 (Proteintech 20824-1-AP), tubulin (Novus NB100-690). Secondary antibodies used were diluted to 1:10000 and incubated at room temperature for 1 hour: Donkey anti-Mouse IgG (H+L) (Invitrogen A32789), Donkey anti-Rabbit IgG (H+L) (Invitrogen A32802). Blots were imaged with the Li-Cor Odyssey CLx infrared imaging system. Raw blots shown in SFigure 7.

### Gas Chromatography-Mass Spectrometry (GC-MS) for MICOS KO

Samples were extracted for metabolites and prepared as previously designed^33,110^. Samples were extracted for metabolites in −80°C 2:2:1 methanol/acetonitrile/water that contained a mixture of nine internal standards (d_4_-citric acid, ^13^C_5_-glutamine, ^13^C_5_-glutamic acid, ^13^C_6_-lysine, ^13^C_5_-methionine, ^13^C_3_-serine, d_4_-succinic acid, ^13^C_11_-tryptophan, d_8_-valine; Cambridge Isotope Laboratories) at a concentration of 1 µg/mL each, at a ratio of 18:1 (extraction solvent:sample volume). Cells were lyophilized overnight before extraction and homogenized with a ceramic bead mill homogenizer after the addition of extraction buffer. Samples were incubated for one hour at −20°C and centrifuged at maximum speed for 10 min. All supernatants were transferred to fresh tubes and pooled quality control (QC) samples were prepared by adding an equal volume of each sample to a fresh 1.5 mL microcentrifuge tube. A speed vac was used to evaporate the pooled QCs, samples, and processing blanks, which were made by adding extraction solvent to microcentrifuge tubes. Derivatives of the dried products were obtained using methoxamine hydrochloride (MOX) and *N*,O- bis(trimethylsilyl)trifluoroacetamide (TMS). Products were rehydrated in 30 μL of 11.4 mg/mL Molybdenum Carbide (MOC) in anhydrous pyridine (VWR), vortexed for 10 minutes, and incubated at 60°C for 1 hour. Then 20 µL of TMS was added, vortexing was repeated for one min, and samples were heated for an hour at 60°C. Samples of 1 µL were analyzed by GC-MS using a Thermo Trace 1300 GC with a TraceGold TG-5SilMS column for GC chromatographic separation. The GC temperature was set at 250°C for the inlet, with the oven temperature at a gradient with 3 min at 80 °C, increasing by 20°C a minute to a final 280°C for the last 8 minutes. The settings for the GC machine were 20:1 split ratio, split flow; 24 μL/min, purge flow; 5 mL/min, carrier mode; constant flow, carrier flow rate: 1.2 mL/min. Between each injection sample, the column was washed three times with pyridine. Metabolites were detected using the Thermo ISQ single quadrupole mass spectrometer, with data acquired from 3.90 to 21.00 min in the EI mode (70 eV) by single-ion monitoring. The profiling of the metabolites was performed using TraceFinder 4.1 with standard verified peaks and retention times. TraceFinder was used to compare metabolite peaks in each sample against an in-house library of standards. For these standards, retention times and fragment ions for each were analyzed, with fragment ions for both the target peak and two confirming ions. For the samples, we identified metabolites that matched both retention times and the three fragment ions. TraceFinder was also used for GC-MS peak integration to obtain peak areas for each metabolite. The profiling of the metabolites was performed using TraceFinder 4.1 with standard verified peaks and retention times. TraceFinder was used to compare metabolite peaks in each sample against an in-house library of standards. TraceFinder was also used for GC-MS peak integration to obtain peak areas for each metabolite. After this analysis, we used previously described protocols^111^ to correct for drift over time by using QC samples run at both the beginning and end of the sequence. The data was then normalized to an internal standard to control for extraction, derivatization, and/or loading effects.

### Liquid Chromatography-Mass Spectrometry (LC-MS) for MICOS KO

Myotubes were dried, rehydrated in 40 µL acetonitrile:water (1:1), and vortexed. For LC-MS, 2 µL of the sample was used with a Thermo Q Exactive hybrid quadrupole Orbitrap mass spectrometer with a Vanquish Flex UHPLC system and a Millipore SeQuant ZIC-pHILIC column (length area = 2.1 × 150 mm, 5 µm particle size) with a ZIC-pHILIC guard column (length area = 20 × 2.1 mm). The mobile phase comprised solvent A (20 mM ammonium carbonate [(NH_4_)_2_CO_3_] and 0.1% ammonium hydroxide [NH_4_OH]) and solvent B (acetonitrile). The mobile phase gradient started at 80% solvent B, decreased to 20% solvent B over 20 min, returned to 80% solvent B in 0.5 min, and was held at 80% for 7 min ^112^. From there, the mass spectrometer was operated in the full-scan, polarity-switching mode for 1 to 20 min, spray voltage set to 3.0 kV, capillary heated at 275°C, and HESI probe heated at 350°C. The sheath gas flow, auxiliary gas flow, and sweep gas flow were 40 units, 15 units, and 1 unit, respectively. We examined an *m*/*z* range of 70–1000, the resolution was set at 70,000, the AGC target at 1 × 10^6^, and the maximum injection time was set to 200 ms ^111^. TraceFinder 4.1 software was used for analysis and metabolites were identified based on an in-house library. Drift was corrected for as described above^111^. Data were normalized and further visualization and analysis were performed on MetaboAnalyst 5.0^113^.

### Analyzing Metabolomic Data for MICOS KO

Metabolomic analysis was performed as described previously^33^ using the web service MetaboAnalyst 5.0 (https://www.metaboanalyst.ca/MetaboAnalyst/ModuleView.xhtml, last accessed on 8 February 2022) that combines machine learning methods and statistics to group data using PCA, heat mapping, metabolite set enrichment analysis, and statistical analysis. One-way ANOVA and Fisher’s LSD multiple comparison test were also used. PCA uses score plots to provide an overview of variance for the principal components. Heatmaps separate hierarchical clusters leading to progressively larger clusters. Clusters are based on similarity using Euclidean distance and Ward’s linkage to minimize the clustering needed. Metabolite Set Enrichment Analysis (MSEA), which determines whether a set of functionally related metabolites is altered, can be used to identify consistent changes across many metabolites with similar roles.

Overrepresentation analysis determines whether a group of compounds is overrepresented in comparison to pure chance and whether a group of metabolites have similar changes. In this analysis, the fold enrichment was calculated by dividing the observed hits by the expected metabolites. Expected number of hits are calculated by MetaboAnalyst 5.0. GraphPad Prism software (La Jolla, CA, USA) was used for statistical analysis with data expressed as mean ± standard deviation, and one-tailed p-values ≤ 0.01 were considered significant.

### Metabolomics for Aged Samples

Methods used previously verified techniques ^114–116^. Frozen tissues were weighed, ground with a liquid nitrogen in a cryomill (Retsch) at 25 Hz for 45 seconds, before extracting tissues 40:40:20 acetonitrile: methanol: water +0.5% FA +15% NH4HCO3 ^116^ with a volume of 40µL solvent per 1mg of tissue, vortexed for 15 seconds, and incubated on dry ice for 10 minutes. Tissue samples were then centrifuged at 16,000 g for 30 minutes. The supernatants were transferred to new Eppendorf tubes and then centrifuged again at 16,000 g for 25 minutes to remove and residual debris before analysis.

Extracts were analyzed within 24 hours by liquid chromatography coupled to a mass spectrometer (LC-MS). The LC–MS method was based on hydrophilic interaction chromatography (HILIC) coupled to the Orbitrap Exploris 240 mass spectrometer (Thermo Scientific) ^114^. The LC separation was performed on a XBridge BEH Amide column (2.1 x 150 mm, 3.5 μm particle size, Waters, Milford, MA). Solvent A is 95%: 5% H2O: acetonitrile with 20 mM ammonium acetate and 20mM ammonium hydroxide, and solvent B is 90%: 10% acetonitrile: H2O with 20 mM ammonium acetate and 20mM ammonium hydroxide. The gradient was 0 min, 90% B; 2 min, 90% B; 3 min, 75% B; 5 min, 75% B; 6 min, 75% B; 7 min, 75% B; 8 min, 70% B; 9 min, 70% B; 10 min, 50% B; 12 min, 50% B; 13 min, 25% B; 14min, 25% B; 16 min, 0% B; 18 min, 0% B; 20 min, 0% B; 21 min, 90% B; 25 min, 90% B. The following parameters were maintained during the LC analysis: flow rate 150 mL/min, column temperature 25 °C, injection volume 5 µL and autosampler temperature was 5 °C. For the detection of metabolites, the mass spectrometer was operated in both negative and positive ion mode. The following parameters were maintained during the MS analysis: resolution of 180,000 at m/z 200, automatic gain control (AGC) target at 3e6, maximum injection time of 30 ms and scan range of m/z 70-1000. Raw LC/MS data were converted to mzXML format using the command line “msconvert” utility ^115^. Data were analyzed via the EL-MAVEN software version 12.

### Lipidomics for Aged Samples

#### Tissue homogenization and extraction for lipids

Tissues were homogenized using a Retsch CryoMill. The homogenate was mixed with 1 mL of Extraction Buffer containing IPA/H2O/Ethyl Acetate (30:10:60, v/v/v) and Avanti Lipidomix Internal Standard (diluted 1:1000) (Avanti Polar Lipids, Inc. Alabaster, AL). Samples were vortexed and transferred to bead mill tubes for homogenization using a VWR Bead Mill at 6000 g for 30 seconds, repeated twice. The samples were then sonicated for 5 minutes and centrifuged at 15,000 g for 5 minutes at 4°C. The upper phase was transferred to a new tube and kept at 4°C. To re-extract the tissues, another 1 mL of Extraction Buffer (30:10:60, v/v/v) was added to the tissue pellet-containing tube. The samples were vortexed, homogenized, sonicated, and centrifuged as described earlier. The supernatants from both extractions were combined, and the organic phase was dried under liquid nitrogen gas.

#### Sample reconstitution for lipids

The dried samples were reconstituted in 300 µL of Solvent A (IPA/ACN/H2O, 45:35:20, v/v/v). After brief vortexing, the samples were sonicated for 7 minutes and centrifuged at 15,000 g for 10 minutes at 4°C. The supernatants were transferred to clean tubes and centrifuged again for 5 minutes at 15,000 g at 4°C to remove any remaining particulates. For LC-MS lipidomic analysis, 60 µL of the sample extracts were transferred to mass spectrometry vials.

#### LC-MS analysis for lipids

Sample analysis was performed within 36 hours after extraction using a Vanquish UHPLC system coupled with an Orbitrap Exploris 240™ mass spectrometer equipped with a H-ESI™ ion source (all Thermo Fisher Scientific). A Waters (Milford, MA) CSH C18 column (1.0 × 150 mm × 1.7 µm particle size) was used. Solvent A consisted of ACN:H2O (60:40; v/v) with 10 mM Ammonium formate and 0.1% formic acid, while solvent B contained IPA:ACN (95:5; v/v) with 10 mM Ammonium formate and 0.1% formic acid. The mobile phase flow rate was set at 0.11 mL/min, and the column temperature was maintained at 65 °C. The gradient for solvent B was as follows: 0 min 15% (B), 0–2 min 30% (B), 2–2.5 min 48% (B), 2.5–11 min 82% (B), 11–11.01 min 99% (B), 11.01–12.95 min 99% (B), 12.95–13 min 15% (B), and 13–15 min 15% (B).

Ion source spray voltages were set at 4,000 V and 3,000 V in positive and negative mode, respectively. Full scan mass spectrometry was conducted with a scan range from 200 to 1000 m/z, and AcquireX mode was utilized with a stepped collision energy of 30% with a 5% spread for fragment ion MS/MS scan.

#### Lipidomics Analysis

After normalization, all data was analyzed in R using the *lipidr* package (Mohamed et al. 2020). All code to analyze data and generate figures can be found here: https://github.com/mphillips67/Lipidomic-Analysis-Young-and-Aged-Mouse-Tissue. Data sets associated with each tissue type we looked at were analyzes independently. Prior to running any analysis, data was log transformed and further processed. For lipids where there were multiple readings across replicates, sets of readings with the highest values were identified and all others were discarded. Next there were several instances (2 to 3) per tissue type were all samples had identical measurements. As this was mostly likely due to technical errors, these entries were also discarded from the data set. Lastly, to be compatible with *lipidr*, lipid names had to be modified to fit a standard “CLS xx:x/yy:y” naming scheme where CLS refers to the abbreviated lipid class and xx:x and yy:y refer to the first and second chains (note: code to generate a conversion key are available through the previously mentioned GitHub repository).

After processing, lipid composition between young and old samples were compared using the “de_analysis” function from *lipidr* with default settings. Here *lipidr* uses moderated t-tests to identify significantly differentiated lipids between samples types within a given tissue type.

Significantly differentiated lipids are those with adjusted p-values < 0.05 (note: p-values were adjusted to correct for multiple comparisons using and FDR procedure) and log fold changes greater than 1 or less than -1. These results were then used to perform a lipid set enrichment analysis using the “lsea” function where entries were ranked by fold change, only classes with at least 4 associated lipids were considered, and 100000 permutations were run. Here the method *lipdr* used is based on the commonly used gene set enrichment analysis approach outlined in Subramanian et al. (2005). Briefly, lipid class and chain length categories are determined from annotations extracted from lipid names in the data set and lipids are ranked by fold change. A permutation algorithm is then used to calculate enrichment scores and p-values for each lipid set. Sets with adjusted p-values < 0.05 were defined as significantly enriched. Lastly, heatmaps were generated for significantly enriched lipid classes using the “plot_heatmap”.

### Data Analysis

All SBF-SEM and TEM data were presented as the mean of at least three independent experiments with similar outcomes. Results were presented as mean ± standard error with individual data points shown. Data with only two groups were analyzed using an unpaired t-test. For nanotunnel quantification, a Mann-Whitney test (unpaired, nonparametric) t-test was performed between two groups. If more than two groups were compared, one-way ANOVA was performed, and significance was assessed using Fisher’s protected least significant difference (LSD) test. GraphPad Prism software package was used for t-tests and ANOVA analyses (La Jolla, CA, USA). For all statistical analyses, *p* < 0.05 indicated a significant difference. Higher degrees of statistical significance (**, ***, ****) were defined as *p* < 0.01, *p* < 0.001, and *p* < 0.0001, respectively.

## ACKNOWLEDGEMENTS

This project was funded by the National Institute of Health (NIH) NIDDK T-32, number DK007563 entitled Multidisciplinary Training in Molecular Endocrinology to Z.V.; National Institute of Health (NIH) NIDDK T-32, number DK007563 entitled Multidisciplinary Training in Molecular Endocrinology to A.C.; Integrated Training in Engineering and Diabetes", Grant Number T32 DK101003; Burroughs Wellcome Fund Postdoctoral Enrichment Program #1022355 to D.S.; The UNCF/ Bristol-Myers Squibb (UNCF/BMS)- E.E. Just Postgraduate Fellowship in Life sciences Fellowship and Burroughs Wellcome Fund/ PDEP #1022376 to H.K.B.; NSF MCB #2011577I to S.A.M.; NSF EES2112556, NSF EES1817282, NSF MCB1955975, and CZI Science Diversity Leadership grant number 2022-253614 from the Chan Zuckerberg Initiative DAF, an advised fund of Silicon Valley Community Foundation to S.D.; The UNCF/Bristol-Myers Squibb E.E. Just Faculty Fund, Career Award at the Scientific Interface (CASI Award) from Burroughs Welcome Fund (BWF) ID # 1021868.01, BWF Ad-hoc Award, NIH Small Research Pilot Subaward to 5R25HL106365-12 from the National Institutes of Health PRIDE Program, DK020593, Vanderbilt Diabetes and Research Training Center for DRTC Alzheimer’s Disease Pilot & Feasibility Program. CZI Science Diversity Leadership grant number 2022- 253529 from the Chan Zuckerberg Initiative DAF, an advised fund of Silicon Valley Community Foundation to A.H.J.; and National Institutes of Health grant HD090061 and the Department of Veterans Affairs Office of Research award I01 BX005352 (to J.G.). Howard Hughes Medical Institute Hanna H. Gray Fellows Program Faculty Phase (Grant# GT15655 awarded to M.R.M); and Burroughs Wellcome Fund PDEP Transition to Faculty (Grant# 1022604 awarded to M.R.M). Additional support was provided by the Vanderbilt Institute for Clinical and Translational Research program supported by the National Center for Research Resources, Grant UL1 RR024975–01, and the National Center for Advancing Translational Sciences, Grant 2 UL1 TR000445–06 and the Cell Imaging Shared Resource. Its contents are solely the responsibility of the authors and do not necessarily represent the official view of the NIH. The funders had no role in study design, data collection and analysis, decision to publish, or preparation of the manuscript. The co-authors would like to acknowledge the Huck Metabolomics Core Facility for use of the UHPLC coupled to the OE240 Mass Spectrometer and staff Drs. Imhoi Koo, Ashley Shay, and Sergei Koshkin for helpful discussions on sample preparation for lipidomics. We would like to thank Mariya Sweetwyne for her advice as an aging expert. We would like to thank Anastasia Berlynn Casper for her contributions to qPCR.

## ETHICAL STANDARDS

The manuscript does not contain clinical studies or patient data.

## CONSENT FOR PUBLICATION

All authors have agreed to the final version of this manuscript.

## DATA AVAILABILITY STATEMENT

The data that support the findings of this study are available from the corresponding author upon reasonable request.

## AUTHOR CONTRIBUTIONS

Z.V., E.G., L.V., J.S., H.K.B., S.A.M., M.A.P., M.R.M., A.H.J., J.A.G., and D.D. conceived and designed research; A.G.M., A.C., L.V., Z.V., T.A.C, B.C.M., J.L., H.K.B., B.R, C.E., D.D., A.C.M., B.C.J., P.P, M.R.M, A.H.J., and J.A.G. performed experiments; J.D., K.N, J.S., E.G., Z.V., J.L., B.R., T.A.C., A.K.R., A.M.Q., V.E., E.G., D.D., A.C.M., B.C.J., P.P, M.R.M., J.A.G., and A.H.J. analyzed data; B.T, K.N, J.S., E.G., Z.V., S.A.M., A.M.Q., V.E., H.K.B., A.C., A.G.M., J.D., M.A.P., M.R.M., D.D., J.A.G., and A.H.J.. interpreted results of experiments and prepared figures; K.N, E.G., Z.V., J.S., S.A.M., L.V., A.G.M., M.A.P., A.K.R., B.C.M., B.T., C.E., A.C., H.K.B., M.R.M., D.D., J.A.G., and A.H.J. drafted manuscript, edited, and revised manuscript; M.R.M., A.H.J., D.D., and J.A.G. approved final version of manuscript.

## (13) SUPPORTING INFORMATION

All data may be requested from the corresponding author.

## Supplementary

**SFigure 1: Distribution of mitochondria quantifications among mice samples.** Mitochondrial volume in (**A**) gastrocnemius, (**B**) soleus, and (**C**) cardiac tissue of 3-months and 2-year murine samples distribution in each individual mouse to represent intra-individual heterogeneity. This is also done for (**D-F**) mitochondrial area, (**G-I**) mitochondrial perimeter, (**J-L**) mitochondrial complexity index, and (**M-O**) mitochondrial sphericity in gastrocnemius, soleus, and cardiac tissue of 3-months and 2-year murine samples.

**SFigure 2: Raw Western blots for gastrocnemius, soleus, and cardiac tissue of tubulin, OPA-1, and mitofilin.**

**SFigure 3: Metabolomics and lipidomic analysis reveal metabolic dysregulation and disruptions in lipid classes and chain lengths with age in gastrocnemius, soleus, and cardiac muscles.** (**A**) Volcano plot of metabolites that were differentially regulated in aged compared to young gastrocnemius, (**B**) soleus, and (**C**) cardiac samples. For our volcano plots, the x-axis represents the median and the y-axis represents the adjusted FDR. (**D**) Enrichment analysis displaying enriched metabolites in aged gastrocnemius, (**E**) soleus, and (**F**) cardiac muscles. (**G**) PCA plot for young and aged gastrocnemius, (**H**) soleus, and (**I**) cardiac samples. (**J**) Lipid class fold-change ratios between young and old tissues in gastrocnemius, (**K**) soleus, and (**L**) cardiac samples. Significantly differentiated lipid classes are those with adjusted p-values < 0.05 (note: p-values were adjusted to correct for multiple comparisons using and FDR procedure) and log fold changes greater than 1 or less than -1. (**M**) Lipid chain-length fold-change ratios between young and old tissues in gastrocnemius, (**N**) soleus, and (**O**) cardiac samples. Significantly differentiated lipid chain lengths are those with adjusted p-values < 0.05 (note: p-values were adjusted to correct for multiple comparisons using and FDR procedure) and log fold changes greater than 1 or less than -1.

## Tables

**Table 1:** Drosophila qPCR flight tissue results of altered MICOS, mitochondrial, and MERCs proteins in relative fold change across the aging process. Fold change represents change in gene expression when comparing population A with population C. Fold change values of >1.0 indicate that the gene is expressed more in the C populations (28- day cycle) than in the A populations (9-day cycle) at day 21. Fold change values of <1.0 indicate that the gene is expressed more in the A populations than in the C populations at day 21. Values of ∼1.0 indicate no difference between the populations. Values deviating from 1.0 by more than 0.20 are highlighted in yellow. This distinction is arbitrary and based on the researchers’ personal interpretation of significance.

| Gene | $\Delta\Delta Ct$ | Fold change |
| --- | --- | --- |
| ATF-4 | -0.781 | **1.718** |
| Drp | -0.781 | **2.980** |
| Marf | -1.764 | **3.397** |
| Opa1 | -2.388 | **5.235** |
| MINOS1 (Mic10) | -0.385 | **1.306** |
| QIL1 (Mic13) | -1.549 | **2.926** |
| CHCHD3/6 (mic19) | -2.279 | **4.853** |
| Dmic60 (mitofilin) | 0.244 | 0.844 |
| APOO (mic26/27) | -1.053 | **2.075** |
| VDAC/Porin | -3.265 | **9.612** |
| IP3R/Itpr | -0.997 | **1.996** |
| FGF21/ bln | -0.140 | 1.102 |
| Ire1 | -1.219 | **2.328** |
| Bip(grp78)/ Hsc 70-3 | -2.259 | **4.786** |
| ATF-6 | 0.072 | 0.951 |
| GRP75/Hsc70-5 | -1.203 | **2.301** |

**Table 2:** qPCR results from *Drosophila* cardiac tissue. Fold change represents change in gene expression when comparing population A with population C. Fold change values of >1.0 indicate that the gene is expressed more in the C populations (28- day cycle) than in the A populations (9-day cycle) at day 21. Fold change values of <1.0 indicate that the gene is expressed more in the A populations than in the C populations at day 21. Values of ∼1.0 indicate no difference between the populations. Values deviating from 1.0 by more than 0.20 are highlighted in yellow. This distinction is arbitrary and based on the researchers’ personal interpretation of significance.

| Gene | $\Delta\Delta Ct$ | Fold change |
| --- | --- | --- |
| ATF-4 | -0.595 | 1.510 |
| Drp | 0.010 | 0.993 |
| Marf | 0.032 | 0.978 |
| Opa1 | -0.428 | 1.346 |
| MINOS1 (Mic10) | -0.452 | 1.368 |
| QIL1 (Mic13) | 0.012 | 0.992 |
| CHCHD3/6 (mic19) | 0.123 | 0.918 |
| Dmic60 (mitofilin) | -1.791 | 3.460 |
| APOO (mic26/27) | -0.012 | 1.008 |
| VDAC/Porin | 0.290 | 0.818 |
| IP3R/Itpr | 0.243 | 0.845 |
| FGF21/ bln | 0.927 | 0.526 |
| Ire1 | 0.221 | 0.858 |
| Bip(grp78)/ Hsc 70-3 | -0.223 | 1.167 |
| ATF-6 | 0.141 | 0.907 |
| GRP75/Hsc70-5 | -0.224 | 1.168 |

## Supplementary Tables

Supplementary Table 1. Lipid classes present in our dataset and associated abbreviations.

Supplementary Table 2. Results for set enrichment analysis, lipid class and chain length, based on lipidomic differentiation between young and old gastrocnemius tissue.

Supplementary Table 3. Results for set enrichment analysis, lipid class and chain length, based on lipidomic differentiation between young and old soleus tissue.

Supplementary Table 4. Results for set enrichment analysis, lipid class and chain length, based on lipidomic differentiation between young and old heart tissue.

