## Supplementary figures and images for "3D Reconstruction of Murine Mitochondria Exhibits Changes in Structure Across Aging Linked to the MICOS Complex"

### Supplemental Figures

#### Supplement Figure 1

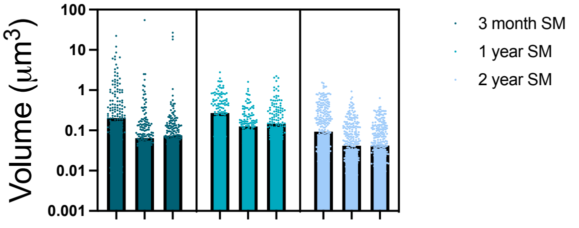


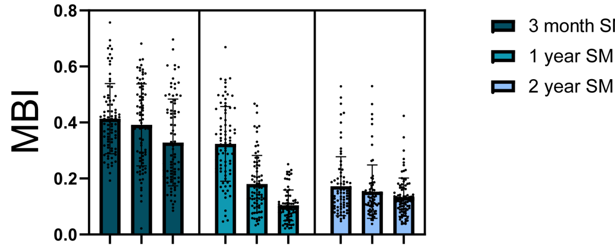


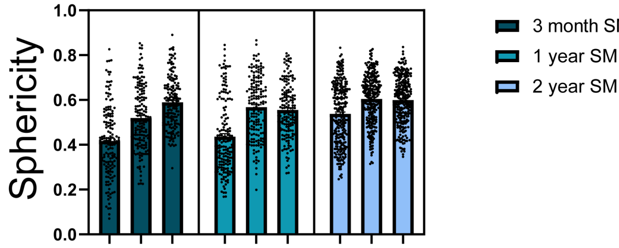


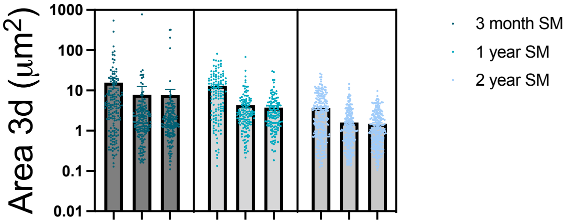


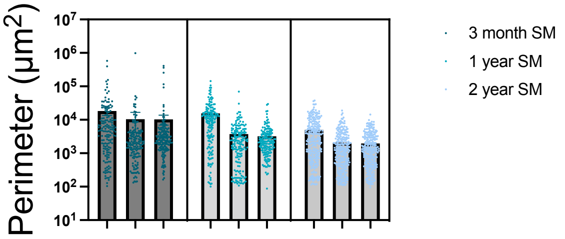

Figure 2. Supplement

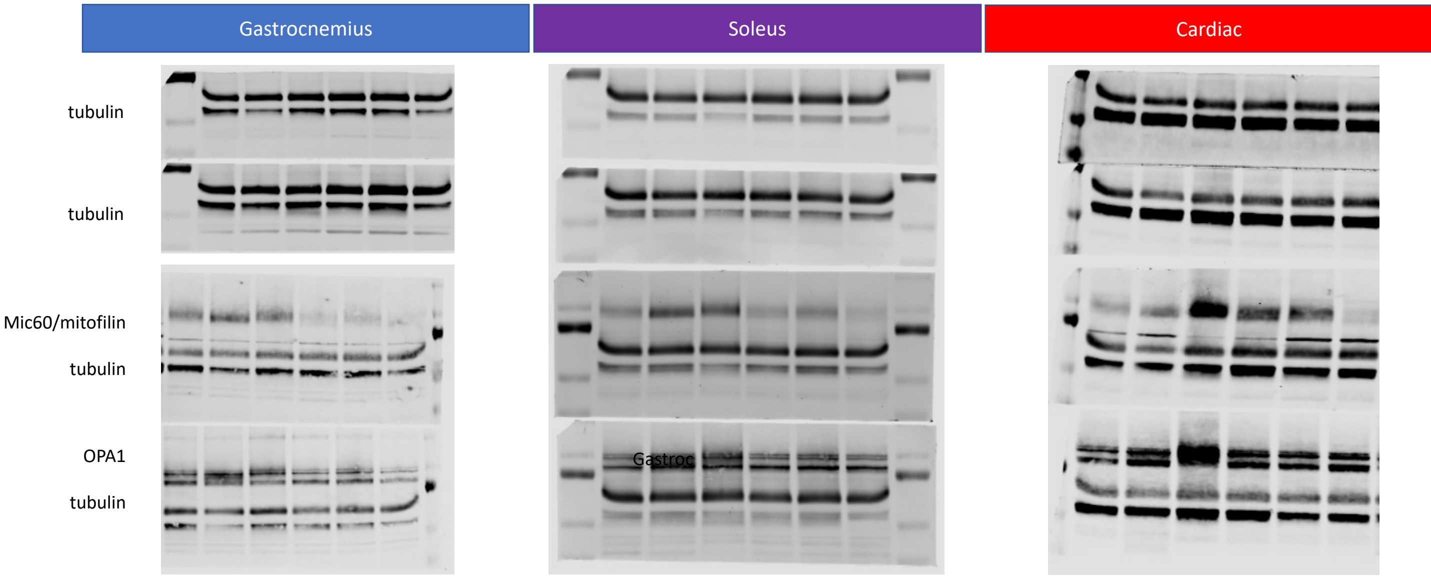

Figure 3 Supplement.

| Gastrocnemius | Soleus | Cardiac |
| --- | --- | --- |

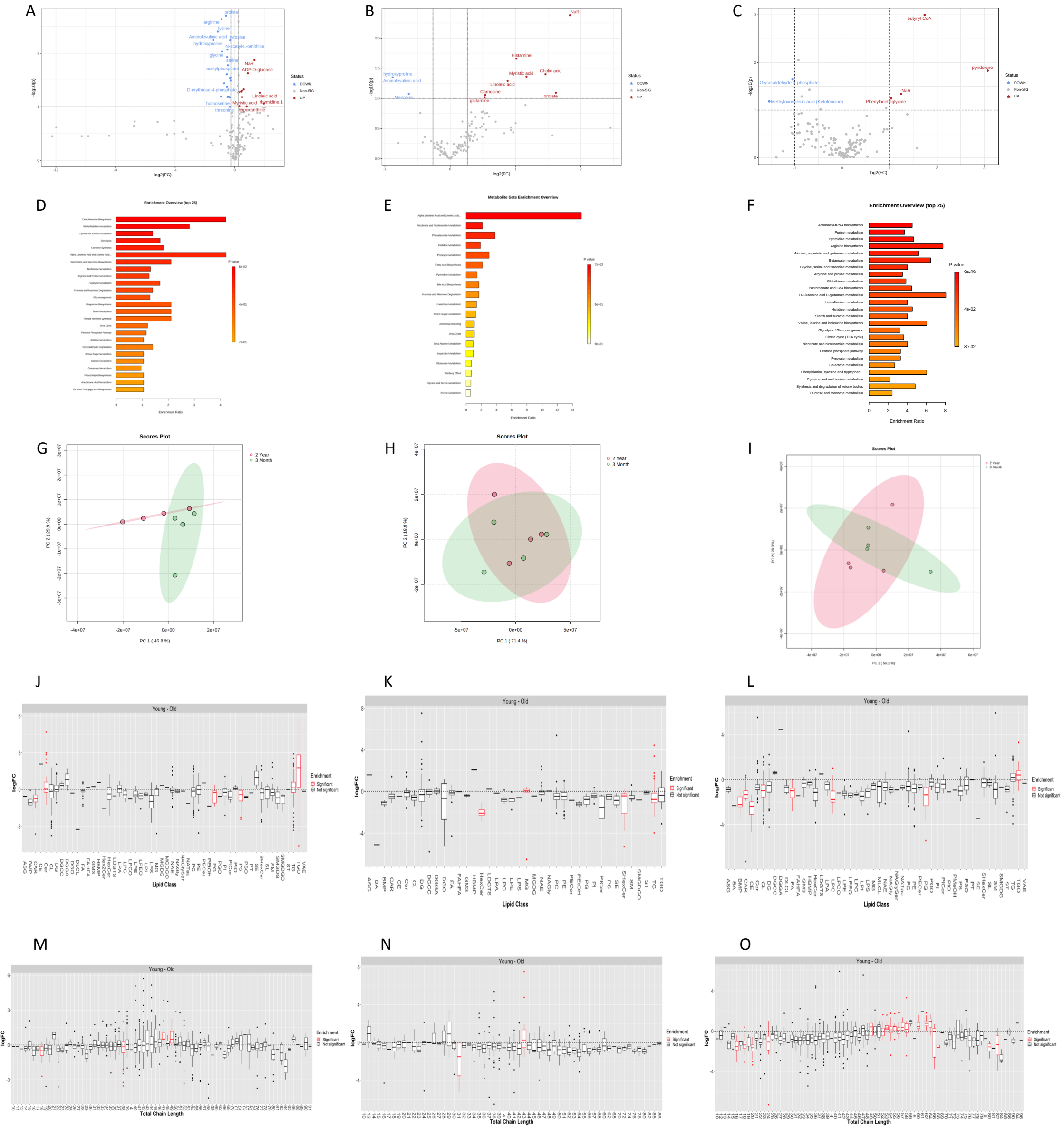
